# A novel PEX14/PEX5 interface links peroxisomal protein import and receptor recycling

**DOI:** 10.1101/2023.08.08.552478

**Authors:** Leonidas Emmanouilidis, Jessica Sehr, Katharina Reglinski, Stefan Gaussmann, David Goricanec, Jonathan Kordon, Filipe Menezes, Dominic Waithe, Philip Hublitz, Verian Bader, Konstanze F. Winklhofer, Martin Jung, Wolfgang Schliebs, Christian Eggeling, Ralf Erdmann, Michael Sattler

## Abstract

Newly synthesized peroxisomal proteins are recognized in the cytosol by the cycling receptor PEX5 and directed to a docking complex comprising PEX14 and PEX13 at the peroxisomal membrane. After cargo translocation, the unloaded PEX5 is recycled in an ATP-dependent manner. Receptor docking involves the WxxxF-motifs in the N-terminal domain (NTD) of PEX5 that are recognized by the N-terminal domain of PEX14. Here, we combine biochemical methods and NMR spectroscopy to identify a novel binding interface between human PEX5 and PEX14. The interaction involves the PEX5 C-terminal cargo-binding TPR domain and a conserved IPSWQI peptide motif in the C-terminal intrinsically disordered region of PEX14. The three-dimensional structure of the PEX14 IPSWQI peptide bound the PEX5 TPR domain, shows the PEX14 interaction is non-overlapping with PTS1 binding to the TPR domain. Notably, PEX14 IPSWQI motif binding to a hinge region in the TPR domain shows a more open supercoil of the TPR fold that resembles the apo conformation in the absence of PTS1 peptide. Mutation of binding site residues in PEX5 or PEX14 leads to a partial protein import defect and decrease of the steady-state-concentration of PEX5. This resembles the mutant phenotype of cells affected in receptor recycling, suggesting a role in this process.

## Introduction

Compartmentalization is a fundamental principle of eukaryotic cells to create micro-environments with specialized metabolic function. Routing of proteins to the correct compartment requires elaborate machineries for protein targeting and translocation that are well established for mitochondria, the nucleus, the ER, and chloroplasts. In contrast, our current knowledge of the molecular mechanism of protein import into peroxisomes are still poorly understood ^1–4^. Peroxisomes are ubiquitous organelles in eukaryotic cells, which can perform diverse functions as a response to the environmental and cellular state. Most common metabolic functions contribute to Reactive Oxygen Species (ROS) homeostasis and lipid metabolism ^5^. Recently, a role for peroxisomes in innate immunity upon viral infection was reported ^6, 7^. The indispensable nature of the peroxisome is manifested by the numerous diseases and disorders linked to malfunction or absence of those organelles ^8^. Peroxisome biogenesis diseases (PBDs) or single enzyme deficiencies may lead to severe health issues. The range of severity varies greatly from a rather mild phenotype as for hyperoxaluria up to 6-months life-expectancy for many peroxisome biogenesis disorders ^9^.

As peroxisomes entirely lack genetic material, all matrix enzymes are imported post-translationally. Peroxins (acronym Pex/PEX for yeast/human proteins) coordinate this routing. Although not all peroxins are conserved among different species, the general cascade of interactions is found in all eukaryotes ^3, 4, 10, 11^. The peroxisomal protein import is characterized by two striking features. First, peroxisomes can import folded, even oligomeric proteins ^12–14^. Second, the import is performed by cycling import receptors, which shuttle between the cytosol and the peroxisomal membrane ^15, 16^. Most of the peroxisomal matrix proteins are targeted to the organelle via peroxisomal targeting signals (PTS1 or PTS2) located either at the N-terminus (PTS2) or at the C-terminus (PTS1). Proteins harboring a PTS1 or PTS2 are recognized and bound in the cytosol by the cycling import receptors Pex5 and Pex7, respectively ^17–20^. Pex5 recognizes PTS1 cargo with its C-terminal tetratrico peptide repeat (TPR) domain. Subsequently, the cargo-loaded import receptors are directed to a docking complex at the peroxisomal membrane ^21, 22^. The docking complex consists of PEX14 and PEX13 (and Pex17 in yeast). PEX13 and PEX14, both have the capability to bind the receptor cargo-complex ^23, 24^. An interaction between the N-terminal domain of PEX14 with the N-terminal intrinsically disordered region of PEX5 plays a pivotal role in peroxisomal protein import, since it mediates the initial binding of the receptor cargo complex to the membrane ^23, 25–28, 29^. Previous studies using black lipid membrane bilayers have identified Pex14 and Pex5 as key components of a minimal pore, especially the PTS-1-pore ^30^. However, it is still not known how the proteins are transported into the peroxisomal lumen. After translocation of PEX5 and the cargo protein, PEX5 becomes ubiquitinylated and recycled through a retrotranslocation complex in an ATP-dependent manner ^31, 32^. Thus, PEX5 is released from the membrane back to the cytosol where it is deubiquitinated and available for another import cycle. Mechanistic and structural details of cargo and PEX14 release from PEX5 remain elusive. Notably, recent studies in yeast provided strong support for a key role of Pex13 for peroxisomal import, suggesting that the peroxisomal pore forms a phase-separated condensate involving the Pex13 YG-rich region, with some features resembling the nuclear pore ^33, 34^, but molecular and structural mechanisms of transport across this pore remain poorly understood.

Interactions between the key components of the peroxisomal pore involve often multivalent protein-protein interactions that involve intrinsically disordered regions in these proteins. Most importantly, the PEX5-PEX14 interaction is mediated by short helical (di)aromatic peptide motifs, *e.g.*, WxxxF motifs, in the N-terminal region of PEX5 that bind with nanomolar affinity to a small helical N-terminal domain (NTD) of PEX14 ^28, 29, 35, 36^ (Fig.1A), which also have recently been shown to mediate weak membrane interactions of PEX5 ^37^. Interestingly, the binding affinity of PEX5 WxxxF motifs with the PEX14 NTD is similar in solution and at the membrane, presumably supported by an enthalpy/entropy compensation ^37^. The PEX14 NTD has also been shown to bind to (di)aromatic peptide motifs in the C-terminal region of tubulin and thus to play a role for protein import and motility of peroxisomes ^38^. Recently, a cryo-EM structural model has been reported for a region of yeast PEX14 comprising the NTD up to the coiled-coil, which is located after the trans-membrane region and expected mediate PEX14 oligomerization ^39^. The only high-resolution structural information for PEX14 is the N-terminal domain, which forms a three-helical bundle and mediates WxxxF-binding ^35, 40^. The trypanosomal PEX14 NTD is also target for small molecule inhibitors inhibiting the PEX14 NTD/PEX WxxxF interaction to block glycosomal import as a novel therapeutic concept to treat trypanosomiases ^41^. The PEX14 NTD represents less than one quarter of the full-length protein. A hydrophobic stretch of amino acids, involved in membrane association, and the coiled-coil domain are predicted from sequence analysis, but high-resolution structural details are elusive ^15^.

**Figure 1.**
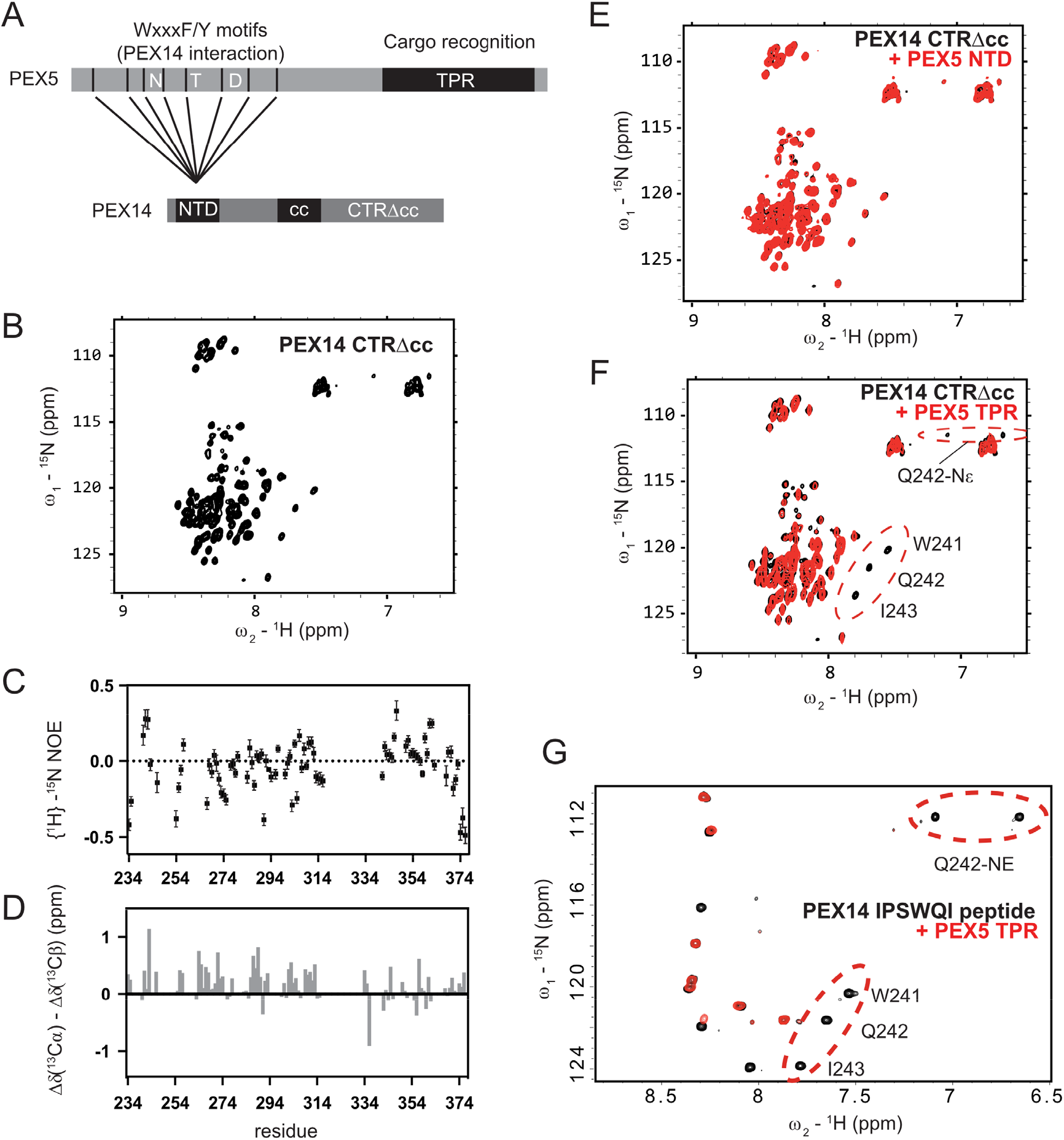
NMR analysis of PEX14 CTRΔcc and its interaction with PEX5. **A)** Schematic representation of the human PEX5 and PEX14 domain organization and their known interactions. NTD: N-terminal domain, TPR: Tetratrico Peptide Repeat, cc: coiled-coil, CTRΔcc: C-terminal domain beyond the coiled-coil domain **B**) ^1^H-^15^N HSQC spectrum of 100 μM PEX14 CTRΔcc in 20 mM phosphate buffer, pH 6.5 and 50 mM NaCl. **C)** {^1^H}-^15^N heteronuclear NOE data and **D)** ^13^C secondary chemical shifts. Error bars in heteronuclear NOE plot correspond to standard deviation of noise in individual spectra. **E, F**) ^1^H-^15^N HSQC spectra of 100 μM PEX14 CTRΔcc (black) and upon addition of equimolar amount of PEX5 NTD (**E**, red) or TPR (**F**, red). **G)** ^1^H-^15^N HSQC spectrum of 150 μM ^15^N-labeled WQI peptide (black) supplemented with 150 μM unlabeled PEX5 TPR.

Here, we present the first structural characterization of the human PEX14 C-terminal region (CTR) beyond the coiled-coil of human PEX14. We show that the PEX14 CTR is intrinsically disordered and highly flexible in solution. Surprisingly, by combining biochemical experiments, NMR spectroscopy and molecular dynamics simulations, we discovered an IPSWQI containing peptide motif in the PEX14 CTR that binds to the cargo-binding C-terminal TPR domain of PEX5. We present the three-dimensional structure of this motif in complex with the PEX5 TPR domain, which shows that the PTS1 peptide binding region and the PEX14 IPSWQI interaction are non-overlapping. The IPSWQI motif binds a flexible hinge region located between the TPR4 and TPR5 repeats in PEX5 with a more open conformation of the TPR supercoil compared to the closed conformation that is observed when the PTS1 peptide is bound^42^. The novel interaction between the C-terminal regions of PEX14 and PEX5 is probed by functional studies in cellular assays, which demonstrate that the interaction is critical for the function of PEX14 and PEX5 in peroxisomal protein import, downstream of receptor docking, presumably during or after cargo release.

## Results

### PEX14 CTRΔcc is intrinsically disordered

To structurally characterize PEX14 CTR, we tested various gene constructs for protein expression and solubility. Inclusion of the putative coiled-coil region to any of the constructs resulted in low expression yields and a heterogeneous population of oligomers. Therefore, we focused on the C-terminal region beyond the coiled-coil domain, named PEX14 CTRΔcc (Fig.1A).

The 25 kDa protein is soluble in aqueous buffers at millimolar concentrations with no detectable oligomerization and was uniformly ^13^C/^15^N isotope-labeled for solution NMR experiments. NMR ^1^H,^15^N correlation spectra show that proton chemical shifts for all backbone amide NMR signals are between 8-8.7 ppm, which indicates that the protein is mostly unstructured (Fig. 1B). In agreement with this, low {^1^H}-^15^N nuclear Overhauser enhancement (NOE) values (Fig. 1C) around zero show that the polypeptide chain is highly flexible in solution at sub-nanosecond time scales. ^13^C secondary chemical shifts, Δδ(^13^Cα) - Δδ(^13^Cβ), indicate the absence of significant secondary structure (Fig. 1D). This demonstrates that the PEX14 C-terminal region beyond the predicted coiled-coiled domain is intrinsically disordered.

### PEX14 CTRΔcc binds in vitro PEX5 TPR domain

To further investigate the molecular functions of the C-terminal region of PEX14 region, we tested whether the PEX14 CTRΔcc can interact with PEX5. We compared ^1^H,^15^N HSQC spectra of ^15^N labeled PEX14 CTRΔcc (Fig. 1B) in the presence of full length PEX5 protein (Supplementary Fig. 1) or in the presence of the PEX5 N-terminal disordered domain (Fig. 1E) or its C-terminal TPR domain (Fig. 1F). Since the PEX5 protein was unlabeled, its presence is not detectable in this heteronuclear NMR correlation experiment. Upon addition of full length PEX5, a number of signals in the spectrum of ^15^N-labeled PEX14 CTRΔcc were broadened beyond detection (Supplementary Fig. 1). The presence of the PEX5 TPR domain causes line-broadening for a number of amide signals in PEX14 (Fig. 1F), corresponding to the same set of signals that are affected upon addition of full length PEX5 (Supplementary Fig. 1). This indicates a potential interaction with PEX5, where the increased molecular weight of PEX14 CTRΔcc in complex with PEX5 is expected to cause line-broadening beyond detection of the NMR signals. Strikingly, upon titration of PEX5 NTD no detectable changes are observed (Fig. 1E). These experiments identified a novel binding interface between PEX14 and PEX5 that involves the C-terminal parts of both proteins. Following NMR chemical shift assignment of the PEX14 NMR signals, we identified the residues that are affected by the PEX5 binding. This analysis revealed a stretch of amino acids in the PEX14 CTRΔcc (residue 234-251, SAPKI<u>PSWQI</u>PVKSPSPS), with a central IPSWQI sequence motif. Since the rest of the PEX14 amide signals are not affected upon addition of the TPR domain, we replaced the protein for further experiments with the 18-mer peptide (residues 234-251), henceforth referred to as IPSWQI peptide. NMR titrations confirm that this peptide binds to the PEX5 TPR domain (Fig. 1G) and affects the same amide signals in PEX5 as observed upon titration of PEX14 CTRΔcc.

We further characterized binding of the PEX14 IPSWQI peptide to ^15^N-labeled PEX5 TPR domain (Fig. 2A) by NMR. Most affected amide signals of the PEX5 TPR domain exhibit fast exchange binding kinetics consistent with a relatively weak interaction (Fig. 2B). In order to map the interaction interface, we assigned the backbone chemical shifts of the PEX5 TPR domain and analyzed the chemical shift perturbations (Fig. 2C). This analysis shows that exclusively the TPR4 and TPR5 motifs of the PEX5 domain (residues 418 to 517) are affected. Notably, this region is distant from the PTS1 binding site of the PEX5 TPR domain. This is illustrated by indicating the PTS1 binding site and the IPSWQI chemical shift perturbations onto the crystal structure of the PEX5 TPR domain ^42^ (Fig. 2D,E).

**Figure. 2.**
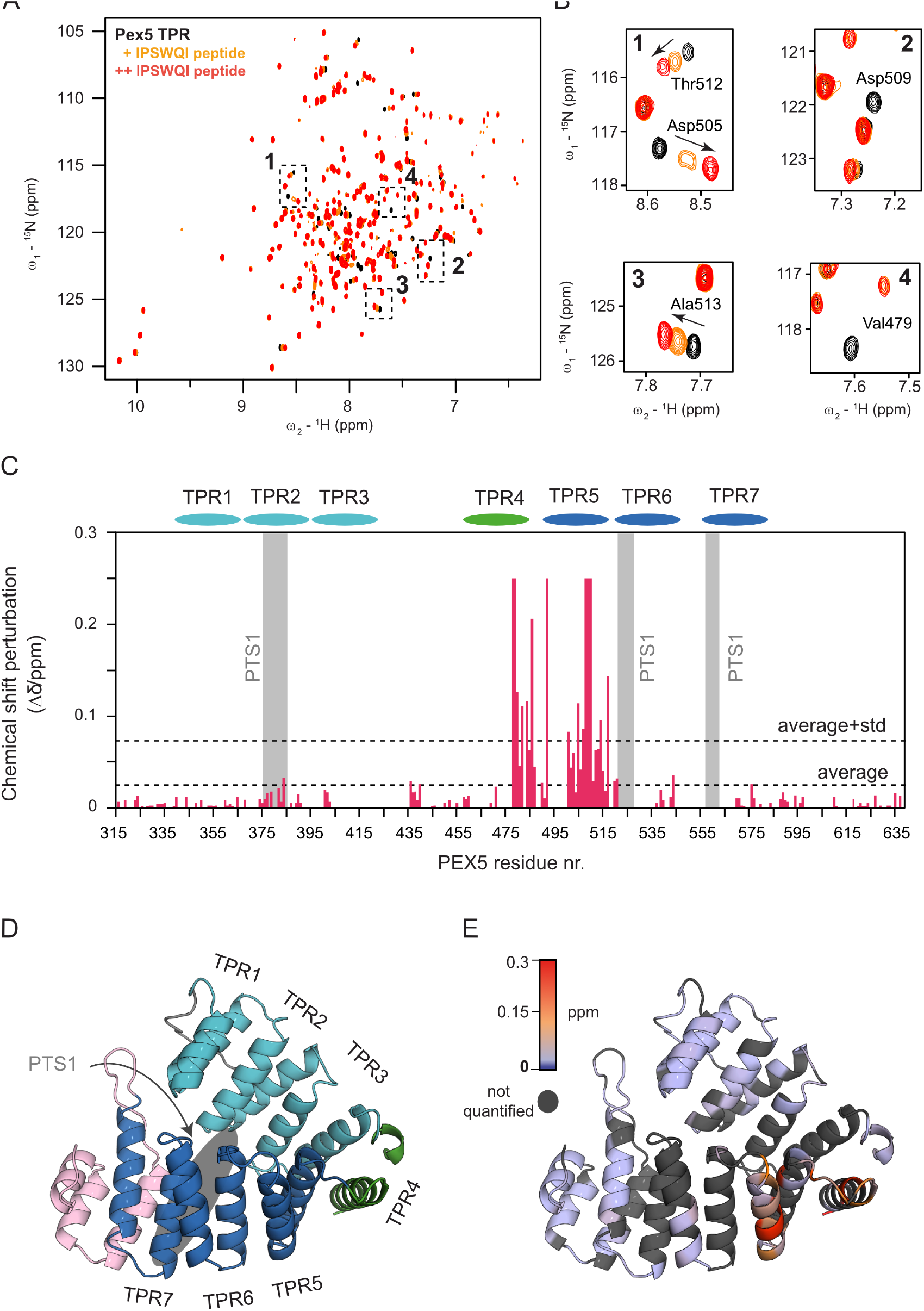
Mapping the PEX14 binding interface with PEX5 TPR. **A)** Overlay of ^1^H-^15^N HSQC NMR spectra of 150 μM PEX5 TPR (black) free and in the presence of 400 μM (orange) and 1800 μM (red) PEX14 IPSWQI peptide, respectively. **B)** Zoomed views show resonances. The number refers to the regions highlighted in A. **C)** Chemical shift perturbations of PEX5 TPR amides upon addition of the PEX14 IPSWQI peptide plotted on the sequence. Line broadened residues beyond detection have set to maximum perturbation of 0.25. PEX5 TPR repeats are indicated on top of the graph. PTS1 binding regions are highlighted with grey boxes. **D)** Crystal structure of the human PEX5 TPR domain (PDB: 2C0M) ^42^. The TPR repeats are annotated and the PTS1 binding cavity is indicated as a grey oval shape. **E)** Cartoon representation of the PEX5 TPR domain colored depending on the extent of chemical shift perturbation. Regions colored in red and blue correspond to most/least affected regions upon WQI peptide binding, respectively. Residues with non-quantified perturbations are colored in gray.

### Structural analysis of the PEX5/PEX14 interaction

We next determined the structure of the PEX5 TPR/PEX14 peptide complex. The conformation of the IPSWQI PEX14 peptide was obtained from transfer NOEs ^43, 44^, exploiting the 36 kDa molecular weight of the PEX5 TPR domain and fast binding kinetics of the peptide ligand. Comparison of NOESY spectra with and without the presence of the TPR domain, revealed NOEs specific to the bound stage. Almost all these transferred NOEs involved the PEX14 tryptophan (W241) side chain. Note, PEX14 and PEX5 residues are denoted by single and triple-letter amino acid codes, respectively. In particular, the aromatic ring shows contacts to the preceding proline (P239) and the following glutamine (Q242) side chains (Supplementary Fig. 2). The bound conformation of the region comprising PEX14 residues 237-245 was calculated using CYANA from a total of 40 short and medium range NOEs (Supplementary Fig. 2; Supplementary Table 1). Analysis of the structural ensemble reveals as a key feature of the bound peptide conformation stacking of the tryptophan ring with P239 and Q242 side chains creating a kink in the peptide backbone (Fig. 3A,B).

**Figure. 3.**
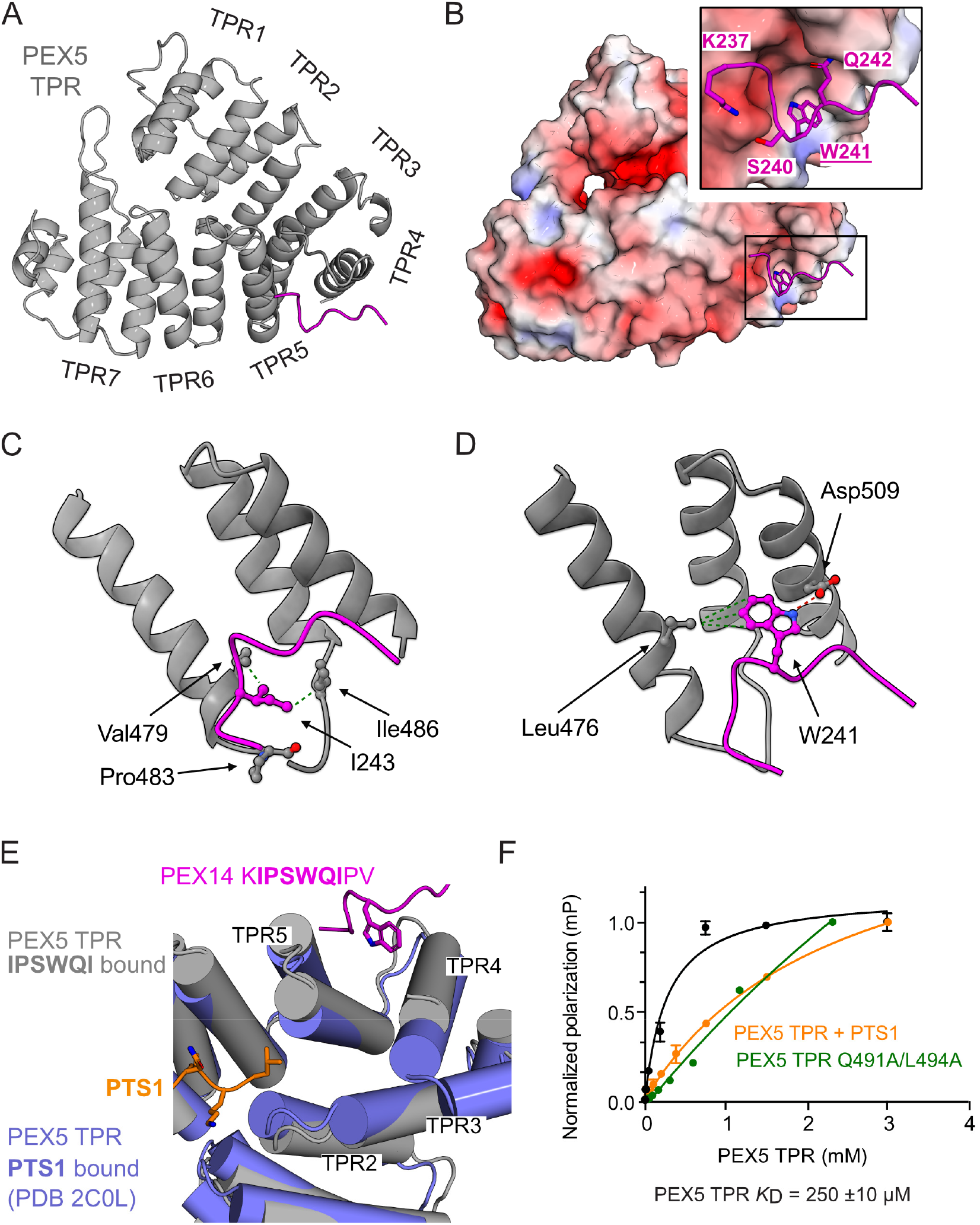
Analysis of the PEX5 TPR - PEX14 IPSWQIP complex structure. **A)** Cartoon of PEX5 (gray) with PEX14 shown as backbone worm (magenta). TPR segments are annotated. **B)** Electrostatic surface representation of PEX5 TPR domain with the IPSWQI peptide shown magenta. The W241 is shown in stick representation. The zoom in into the binding site shows the fit of W241 and electrostatic and polar coordination of the residues K237, S240 and Q242. **C, D)** Representation of the interaction networks of I243 and W241 with PEX5 residues. Aliphatic and hydrophilic interactions are indicated as green and red dashed lines respectively. **E)** Overlay of the PEX5 TPR domain (blue) bound to mSCP2 PTS1 peptide (orange) and the TPR domain bound to the PEX14 IPSWQIP peptide (magenta). The comparison shows a structural rearrangement between the PTS1 and the WQI bound form. **F)** Affinity measurement of N-terminal FITC conjugated SAPK**IPSWQI**PVKSPSPS PEX14 peptide for PEX5 TPR wild type, Q491A/L494A or preloaded with PTS1 peptide (YQSKL) using fluorescence polarization assay. The fluorescently labeled peptide concentration is constant at 20 nM.

^13^C isotope-filtered/edited NOESY experiments ^45^ were measured on the complex to obtain intermolecular contacts. Signal overlap of the methyl region was overcome by selectively ^1^H,^13^C methyl-labeling (δ_1_-methyl-isoleucine, methyl-leucine and methyl-valine) ^46^. This isotope-labeling scheme, together with ^1^H,^13^C HMQC NMR titrations enabled the chemical shift assignment of sidechain methyl groups in the PEX5 TPR domain that are affected upon binding of PEX14 (Supplementary Fig. 3A, B), in particular PEX5 Ile486, Leu476, and Val479 gave rise to intermolecular NOEs (Supplementary Fig. 3C). All three side chains show intermolecular NOEs to protons corresponding to aliphatic protons of the peptide (Supplementary Fig. 3C), suggesting mainly hydrophobic interactions. Additionally, the Leu476 methyl groups show intermolecular NOEs with aromatic protons of the peptide, which are unambiguously assigned to the side chain of PEX14 W241 (Supplementary Fig. 3C).

Notably, the backbone of the peptide is also in close proximity to Ile486 methyl group. In total 10 intermolecular NOEs were identified (Supplementary Table 2). Based on these NOEs and chemical shift perturbations observed for PEX5 and PEX14 distance restraints were derived and used for semi-rigid body docking calculations using HADDOCK2.4^47–49^ (see methods for details, and Supplementary Table 3 and 4 for structural statistics).

Analysis of the 10 lowest energy structures of the main cluster 1 (Supplementary Fig. 4A) shows good convergence with an overall root mean square Cα coordinate deviation (RMSD) of 0.7 Å. Further statistics and binding energetics can be found in Supplementary Table 3, 4 and Supplementary Fig. 4B. To evaluate the stability of the structures, molecular dynamics (MD) simulations were run on the best scored structure in explicit water for 600 ns simulation time using GROMACS ^50^. The coordinate RMSD calculated for each time frame of the simulation time stays constant indicating a stable protein-peptide complex (Supplementary Fig. 5A). Structural analysis shows that the PEX14 IPSWQI peptide binds mainly to a hydrophobic interface located between TPR4 and TPR5 of the PEX5 TPR domain (Fig. 3A,B), which comprises the aliphatic residues PEX5 Leu476, Val479, Leu494, Phe498, and Cys510. This interface provides a shallow hydrophobic groove accommodating the bulky sidechain of PEX14 W241 (Fig. 3B, Supplementary Fig. 4C) with potential stacking with the PEX5 Phe498 aromatic side chain. Notably, this region is surrounded by further hydrophobic residues, PEX5 Pro482 and Ile486 and the aliphatic sidechains from Arg480 and Lys506, which enlarge the hydrophobic area and accommodate PEX14 I238, P239, I243 and P244. Further stabilization is mediated by electrostatic and polar interactions involving PEX5 Asp509 with PEX14 K237 and W241 (Fig. 3B, Supplementary Fig. 4C), consistent with our MD simulations (Supplementary Fig. 5B-D). PEX5 Pro483 holds PEX14 I243 in the hydrophobic pocket surrounded by Val479 and Ile486 (Fig. 3C). Asp 509 helps to keep W241 close to Leu476 via hydrogen bonding (Fig. 3D). We further observed sidechain correlations between Gln491 and Q242 and between Leu494 and W242 (Supplementary Fig. 5E, F), consistent with line broadening observed in the NMR spectra for residues in close proximity (PEX5 Gln491, Leu494 and Asp509.

To further validate the observed mode of interaction, we determined binding affinities of the wildtype PEX5 TPR domain and a double mutant PEX5 TPR Q491A/L494A (PEX5(2A), which affects two residues at the bottom of the PEX14 binding surface, with the PEX14 IPSWQI peptide using a fluorescence polarization assay. The wildtype PEX5 TPR domain binds with a dissociation constant *K*_D_ = 250 μM, while the interaction of the mutant PEX5 TPR domain is strongly reduced (Fig. 3F). Strikingly, in the presence of PTS1-peptide, *i.e.*, representing the cargo-loaded PEX5 conformation, the affinity for PEX14 is strongly reduced compared to the unloaded PEX5 TPR domain. Thus, the PEX14 IPSWQI peptide preferentially binds to cargo-unloaded PEX5. The PEX5(2A) mutant behaved like cargo-loaded PEX5, it clearly lost its capability to interact with PEX14 IPSWQI peptide *in vitro* and is used to assess the functional role of the new PEX5/PEX14 interface *in vivo*.

In this context it is noteworthy, that previous structural analysis of the free PEX5 TPR and complexed with mSCP2, a PTS1 cargo protein, revealed a rearrangement of the PEX5 TPR segments TPR2, TPR3, TPR4, and TPR5, tilting the helices α4B (TPR4) and α5A (TPR5) away from each other. This changes the overall supercoil of the TPR helicase toward a more closed conformation, caused by interactions of the PTS1 peptide with the TPR2 and TPR3^42, 51^ (Supplementary Fig. 4D). Strikingly, this structural rearrangement is turned in the opposite direction to a more open supercoil upon binding of the PEX14 IPSWQI peptide, resembling the apo (PTS1-unbound) conformation (Fig. 3E, Supplementary Fig. 4D). This conformational change of the TPR domain is observed in the structural ensemble of the PEX IPSWQI-bound TPR domains with coordinate RMSD of 0.7 Å (Supplementary Table 4).

To analyze the requirements for PEX5 binding, we probed the contributions of amino acid residues in PEX14 that are critical for the PEX5-TPR interaction by using a mutational peptide scan. To this end, a peptide library representing one or more substitutions within the PEX14 IPSWQI-sequence were analyzed by ligand blotting for their ability to bind the PEX5 TPR domain *in vitro*. Amino acid changes for several residues in the PEX14 ^238^IPSWQI^243^ motif did result in a significant decrease of the binding (Supplementary Fig. 6A). For example, exchange of I238 and W241 to other amino acids impaired the binding, so did the replacement of Gln242 or I243 by aspartate. These observations are fully consistent with the NMR and structural data. A striking effect is also observed by the replacement of three or more residues by alanine, which did result in strongly reduced affinity (Supplementary Fig. 6B). The lowest binding is seen upon replacement of five or six of the residues with alanine.

### Mutation of the PEX14 residues involved in PEX5 binding results in a partial import defect

To investigate the relevance of the identified interaction site for PEX14 function, we analyzed the complementing activity of a PEX14 variant in which the motif is substituted by six alanines (PEX14(6A)) in a cell-based model. To this end, we used previously CRISPR/Cas9 generated PEX14-deleted HEK-293 cells ^52^. Due to the PEX14 deletion, these cells lost their ability to import matrix proteins into peroxisomes. The PEX14-deficient cells and control HEK cells were transfected with a bicistronic vector encoding 1) eGFP-SKL as a marker for peroxisomal import and 2) the wild-type PEX14 or PEX14(6A). Rescue or no rescue of the import defect was monitored by structured illumination microscopy (SIM) on fixed HEK-293 cells after possible import. Indicative for proper peroxisomal protein import, the expression of eGFP-SKL alone in wild-type HEK cells did result in a congruent punctuate pattern colocalizing with PMP70, a peroxisomal marker protein (Fig. 4A, top). In contrast, the PEX14-deficient HEK cells show a diffuse cytosolic or nuclear labelling (Fig. 4A, second row). When PEX14 was co-expressed, most cells showed complementation of the import defect 48 hours after transfection, indicated by the congruent fluorescence pattern of the eGFP-SKL and the peroxisomal marker and the mostly disappeared mislocalization of the eGFP-SKL (Fig.4A, third row). Also, when PEX14(6A) was co-expressed, some cells showed a peroxisomal localization eGFL-SKL, however, after 48 hours most cells still displayed the mutant phenotype, characterized by the mislocalization of the peroxisomal reporter protein (Fig. 4A, bottom).

**Figure 4:**
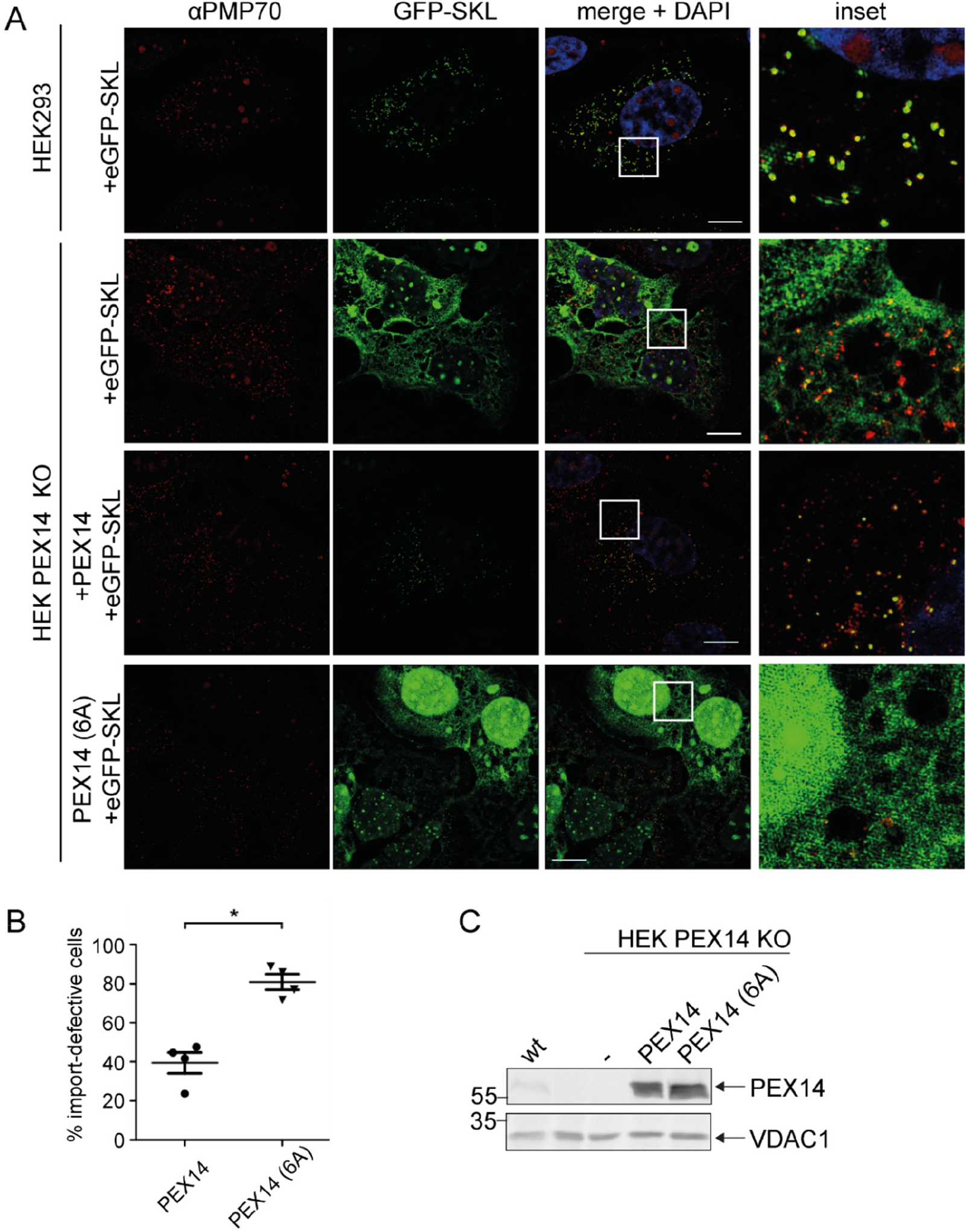
The PEX14 IPSWQI sequence plays a critical role in peroxisomal import of PTS1 proteins. Expression of PEX14(6A) carrying a substitution of the IPSWQI-sequence by AAAAAA does not rescue the peroxisomal import defect as efficient as wildtype PEX14. Both PEX14 constructs were expressed in PEX14-deleted HEK cells from bicistronic expression vectors coding for the indicated PEX14 variants and the eGFP-SKL fusion protein as a marker for PTS1 matrix protein import. **A)** Peroxisomes were labelled by immunofluorescence microscopy of the peroxisomal membrane marker PMP70 and fluorescence images of representative fixed cells were obtained by structured illumination microscopy (SIM) 48 hours after transfection. Peroxisomal co-localization of eGFP-SKL indicates functionality of peroxisomal import. Mislocalization of the reporter to the cytosol and/or the nucleus indicative for a defective import into peroxisomes as typical for non-complemented PEX14-deficient cells. Scale bars: 10 µm. **B)** To quantify the efficacy of the restoration of peroxisomal PTS1-import, ∼100 cells were categorized for each experiment by the localization of the PTS1-import marker eGFP-SKL 48 hours after transfection. Import-deficient cells show an increased cytosolic and nuclear eGFP-SKL fluorescence in comparison to wildtype HEK cells. Values were obtained from four independent experiments and p-values were determined by Mann-Whitney test (*p<0.05). Error bars are shown as mean with SD. **C)** The expression level and stability of the PEX14 variants was checked by immunoblot analysis of whole cell lysates of HEK, HEK PEX14 KO and HEK PEX14 KO cells transfected with a vector coding for PEX14, PEX14(6A), as well as eGFP-SKL using specific polyclonal anti PEX14. An empty vector control (-) showed the specificity of the antibody. Immunoblotting with antibodies against the mitochondrial protein VDAC1 served as a loading control.

To quantify the complementation capability of the mutated PEX14, 100 cells of each condition out of four independent transfections were analyzed for cytosolic and nuclear eGFP fluorescence. Whereas 80% of the cells expressing the mutated PEX14(6A) variant showed a significant mislocalization of the peroxisomal marker protein, only 40% of the PEX14 expressing cells exhibited a diffuse nuclear and cytosolic background staining (Fig. 4B). However, besides the prominent mislocalization, many of the PEX14(6A)-transfected mutant cells also showed targeting of the eGFP-SKL to some peroxisomes, indicative for a partial complementation. In fact, after some time the import defect became less pronounced.

However, the efficiency of this import is impaired even after 5 days as indicated by a lower intensity of the peroxisomal eGFP-SKL signal (Fig. 5). Taken together, our results suggest that the C-terminal PEX5 binding site of PEX14 is not essential for PTS1-dependent import *per se* but that the new binding interface is critical for the efficacy of this process. Immunoblot analyses revealed that the steady-state level of both PEX14 variants is essentially the same (Fig. 4C). Thus, the observed differences in complementing activities are not due to different expression levels or increased degradation of PEX14(6A).

**Figure 5:**
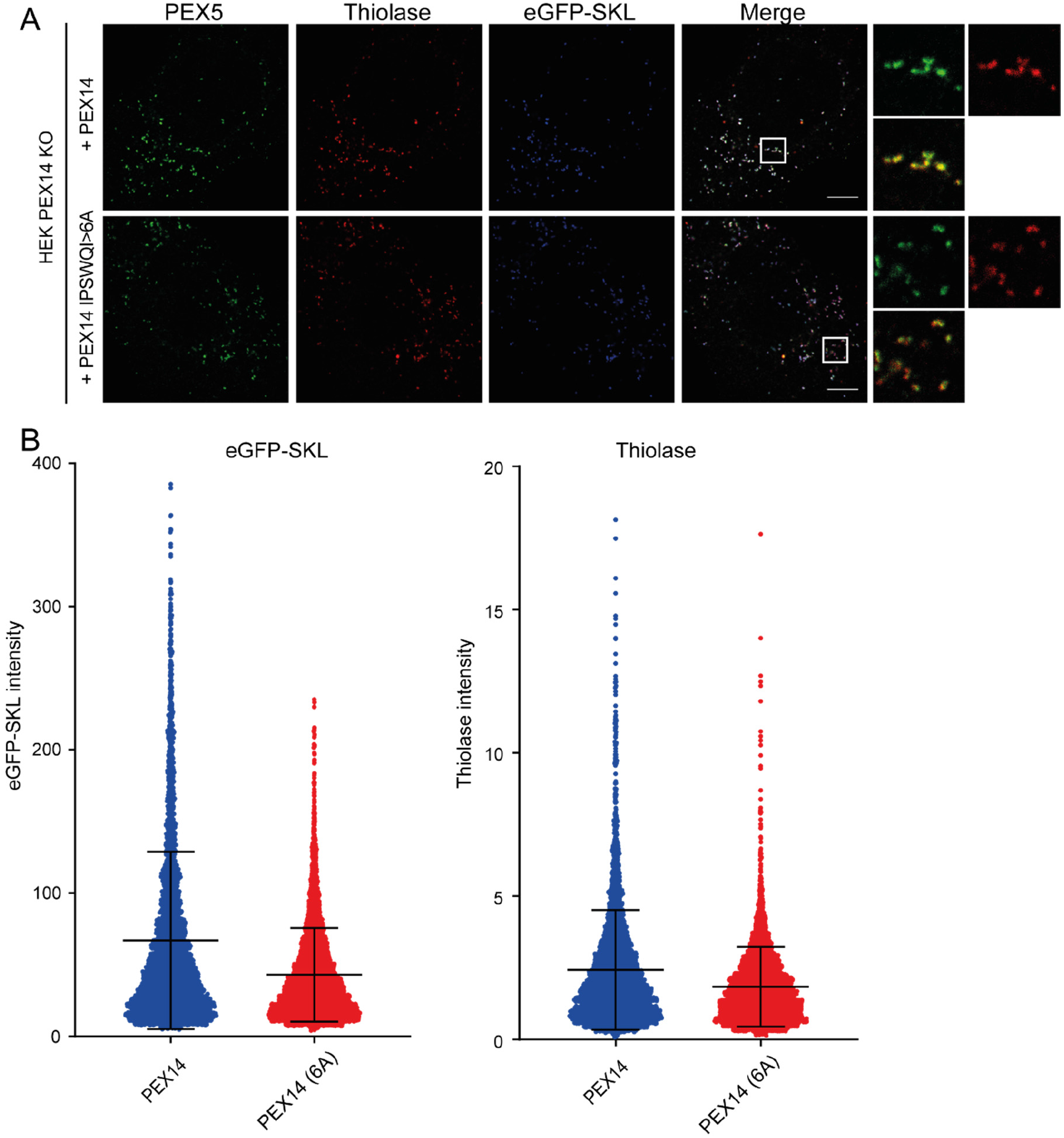
Disruption of the PEX14 interaction with the PEX5 TPR domain has a moderate effect on the PTS2 import pathway. **A)**: Representative super-resolution STED microscopy images of HEK-293 PEX14 KO cells, transfected with a bicistronic vector expressing eGFP-SKL (blue, always confocal) and PEX14 or PEX14(6A), as indicated. The cells were immunolabeled with a monoclonal antibody against membrane-bound PEX5 (green) and polyclonal anti-Thiolase antibodies (red). 4×4 µm intakes are shown to highlight the details of protein distribution at the peroxisomal membrane. In these intakes, brightness and contrast are adjusted to highlight the morphology of the stained proteins. **B)** Images shown in (A) were used for intensity analysis of the PTS2 protein Thiolase. Here the signal for the Thiolase staining was measured in peroxisomal regions of interest, indicated by the eGFP-SKL signal. In the scatter plots, the mean with standard deviation are indicated. The intensity of Thiolase in the cells expressing wild type PEX14 is higher than in the cells expressing the PEX14(6A) variant (p value <0.0001, unpaired T-test).

To test whether the PEX14 IPSWQI motif has an influence on the import of PTS2 proteins, we complemented HEK-293 PEX14 KO cells with wildtype PEX14 or PEX14(6A). In both cases, the PTS2 protein Thiolase was labeled with a specific antibody and the intensity of the labeled Thiolase was measured in images taken with super-resolution STED microscopy on fixed HEK cells after possible import (Fig. 5A). We found that the impact of the IPSWQI motif on the import of the PTS2 protein had a small, but significant effect, which was however much less striking than its effect on the PTS1 protein eGFP-SKL (Fig. 5B).

### Mutagenesis of the PEX5 binding site for PEX14 decreases stability of the PTS1 receptor

The partial import defect caused by disruption of the novel PEX5/PEX14 interface was confirmed by site-directed mutagenesis of the PEX14-IPSWQI core residues to poly A. Next, we investigated whether mutations in the PEX14-binding site of PEX5 show an effect on the activity of the protein in peroxisomal protein import. Our *in vitro* binding studies revealed an essential contribution of PEX5 Q491 and L494 to the interaction with PEX14. As shown by *in vitro* binding assays, substitution of PEX5 amino acids Q491 and L494 by alanine (PEX5L(2A)) reduced PEX14 binding to PEX5 (Fig. 3F).

To analyze the function of PEX5L(2A), harboring Q491A and L494A substitutions, we tested its capability to restore the import defect of PEX5-deficient cell lines ^52^. Using SIM fluorescence imaging on fixed cells the complementation analyses of two independent PEX5 deficient cell lines, PEX5 deleted HEK293 cells (Fig. 6) and Zellweger patient-derived fibroblasts (Supplementary Fig. 7) revealed that the mutated PEX5 protein could not efficiently complement the mutant phenotype of both PEX5-deficient cell lines. This is indicated by the mislocalization of the peroxisomal reporter GFP-SKL to the cytosol and the nucleus (Fig. 6A, lower panel) in comparison to the cell lines complemented with the wild-type PEX5 (Fig. 6A, third panel). In this aspect, PEX5L(2A) seems to cause a similar partial import defect of eGFP-SKL as found with PEX14(6A) (Fig. 4).

**Figure 6:**
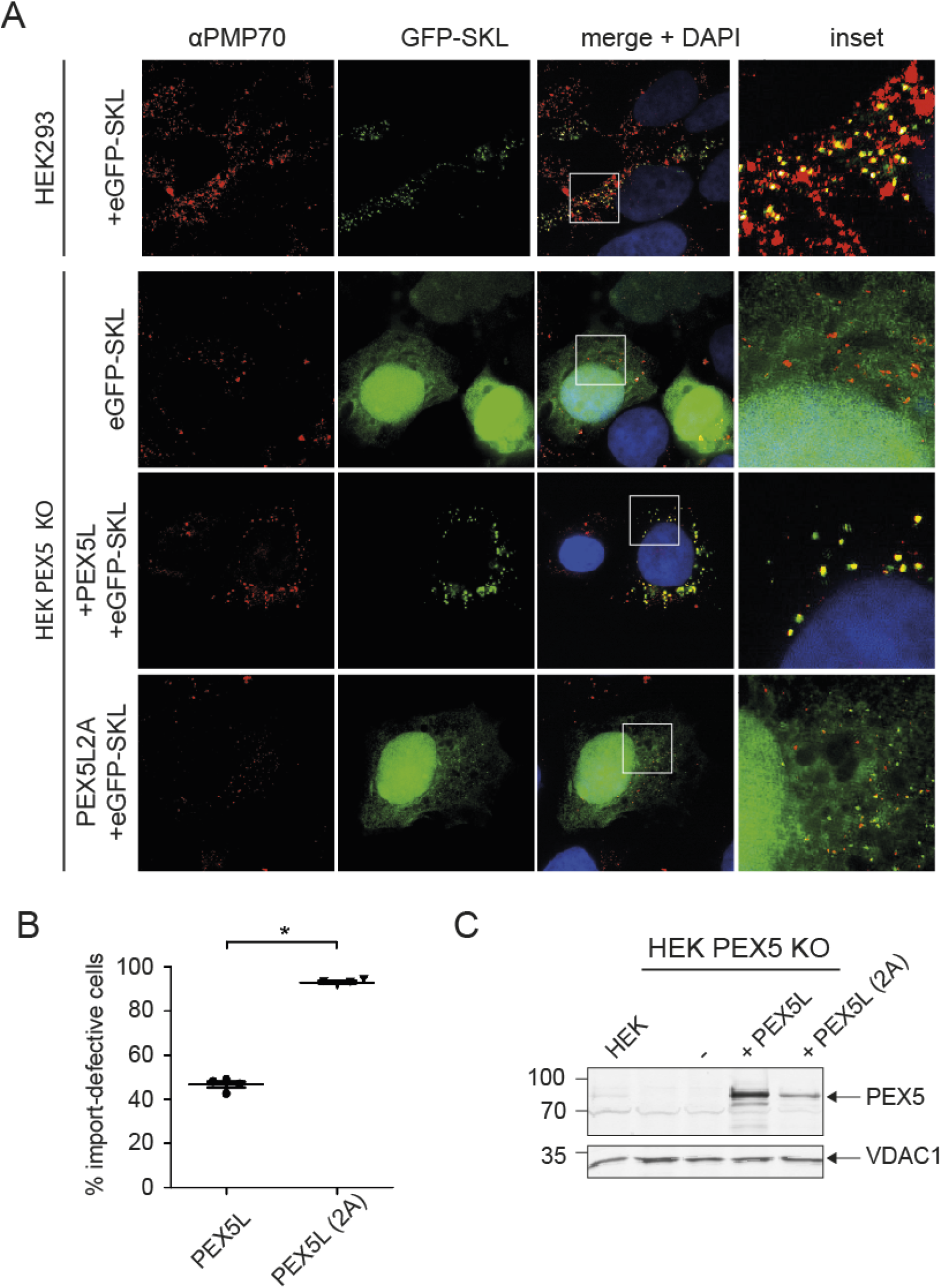
Mutation of the PEX5 binding site for the PEX14 CTR affects peroxisomal protein import and decreases stability of the PTS1 receptor. **A)** The import deficiency of HEK PEX5 knockout cells could be restored by expression of PEX5 and, with lower efficacy, of PEX5L(2A) of which amino acids Q491 and L494 were substituted by alanine. Both constructs were expressed from a bicistronic expression vector, coding for the full-length PEX5 construct (long isoform PEX5L) as indicated and eGFP-SKL as a fluorescent PTS1 matrix protein. Peroxisomes were labeled red by immunofluorescence microscopy using the membrane protein PMP70 as a marker and images on fixed cells were obtained by SIM. Introduction of PEX5L leads to a green punctate pattern, colocalizing with the PMP70. Whereas expression of PEX5 PEX5L(2A) causes a mainly cytosolic distribution of eGFP-SKL. Scale bars: 10 µm. **B**) To quantify the complementation of peroxisomal PTS1-import, for each experiment ∼100 cells were categorized by the localization of the PTS1-import marker eGFP-SKL. Import deficient cells show an increased cytosolic and often nuclear fluorescent labelling in comparison to wildtype HEK cells, with plasmid-encoded expression of eGFP-SKL. Values were obtained from four independent experiments and p-values were determined by Mann-Whitney test (*p≤0.05; **p≤0.01; und ***p≤0.001). Error bars are shown as mean with SD. **C**) The expression level and stability of the PEX5 variants were checked by separating whole cell lysates via SDS-PAGE and analyzed on a immunoblotting using specific polyclonal PEX5 antibody. An empty vector control (pIRES2) shows the specificity of the antibody and anti VDAC1 antibodies were used for loading control.

The quantitative analysis revealed that 50% of the PEX5-deficient cells transfected with wildtype PEX5 plasmid exhibited complete import into peroxisomes. In contrast, almost all transfected cells, expressing the Q491A/L494A variant (PEX5L(2A)) showed mistargeting of GFP-SKL to cytosol and nucleus and only partial labelling of peroxisomes (Fig. 6B). Immunoblot analysis of PEX5 and PEX5L(2A) indicated that the steady-state levels of both PEX5 variants differ. The abundance of PEX5L(2A) in cell lysates was 5-10 fold lower when compared with PEX5 (Fig 6; Supplementary Fig. 8). Analysis of the control (VDAC1) shows that this difference is not caused by different loadings (Fig. 6C). Also, eGFP-SKL and PEX5 variants are translated from the same bicistronic mRNA, thus equal amounts of eGFP-SKL in transfected cell lines exclude the possibility that the reduced steady state level is due to a lower transfection rate (Supplementary Fig. 7). Therefore, we speculate that the partial import defect caused by disruption of the PEX5/PEX14 interface correlates with a reduced stability of the PTS1 receptor.

### Mutational analysis of the interface between the PEX5 and PEX14 CTR leads to an accumulation of the PTS1 receptor at the peroxisomal membrane

To investigate if there is an effect of the PEX14(6A) on the recruitment of endogenous PEX5 to the peroxisomal membrane, we carried out an intensity analysis on images of fixed HEK cells (after possible import) by super-resolution STED microscopy. To this end, wildtype PEX14 and PEX14(6A) were expressed in the PEX14 KO cell line and the intensity of immunolabeled PEX5 and PEX14 as well as the signal of the eGFP-SKL at the peroxisomes were measured. To exclude side effects of not fully complemented cells, we imaged the cells five days after transfection. The import of peroxisomal matrix proteins was restored by both PEX14 as well as PEX14(6A) as indicated by the congruent punctuate fluorescence pattern of the eGFP-SKL, PEX5 and PEX14 (Fig. 7A). However, the efficiency of this import was impaired as indicated by a lower intensity of the eGFP-SKL signal (Fig. 5B). Immunolabeling of PEX5 was carried out by using a monoclonal antibody which specifically detects the membrane-bound form of the PTS1-receptor ^53, 54^. In the PEX14(6A)-transfected mutant cells, the intensity of both peroxisome-associated PEX5 and PEX14 decreased (Fig. 7B). Even though there was significantly less peroxisome-associated PEX14(6A), the amount of PEX5 at the membrane was not proportionally affected. Accordingly, the ratio of fluorescence intensities of PEX5 to PEX14 was slightly but significantly increased in PEX14(6A) compared to PEX14-transfected cells, indicating that relatively more PEX5 binds to PEX14(6A) at the peroxisomal membrane (Fig. 7B).

**Figure 7:**
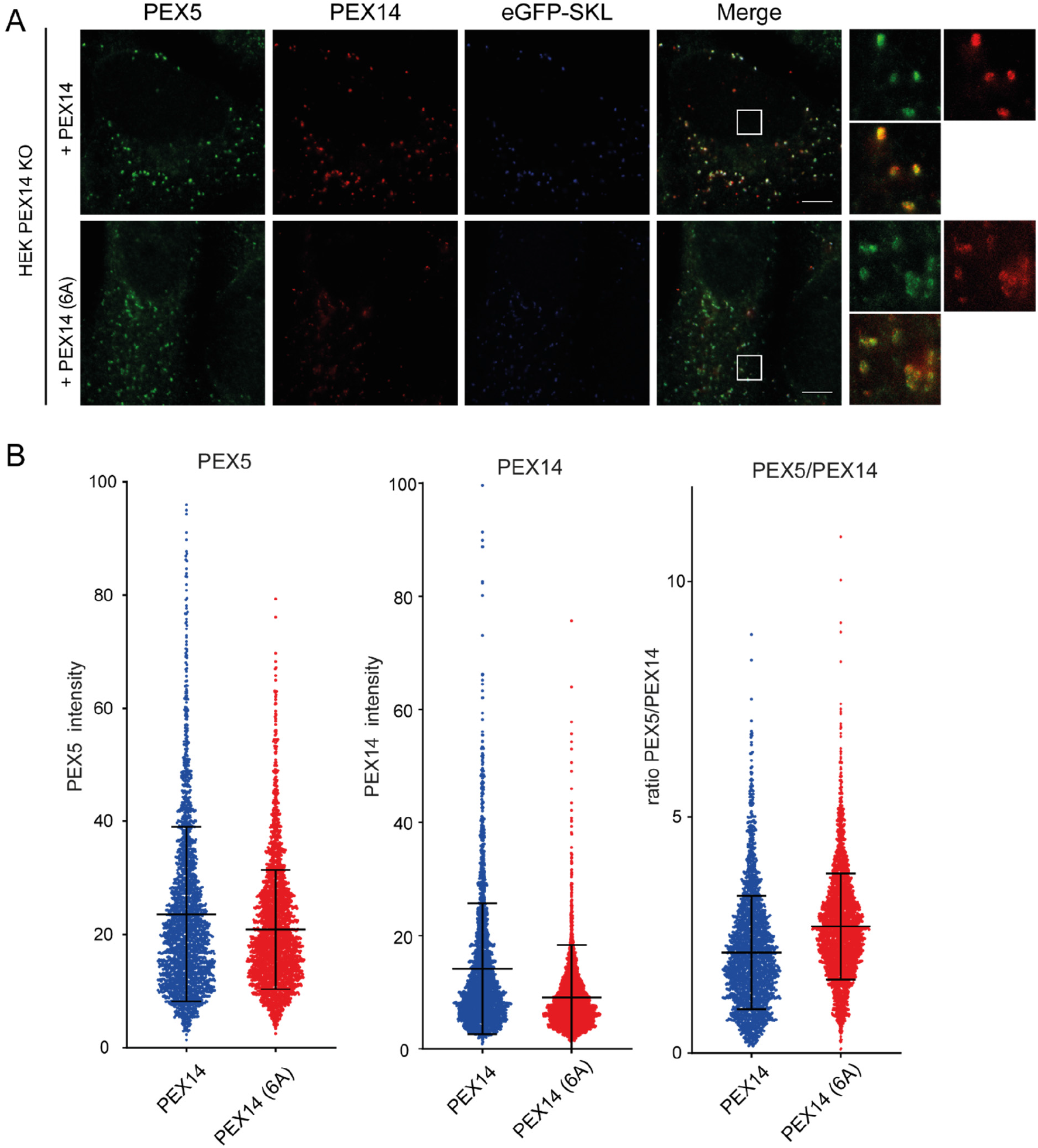
Mutagenesis of the IPSWQI sequence of PEX14 increases the concentration of PEX5 at peroxisomal membranes. **A)** Representative STED microscopy images of HEK-293 PEX14 KO cells, transfected with a dual plasmid expressing eGFP-SKL (blue, always confocal) and a PEX14 or PEX14(6A), as indicated. The cells were immunolabeled for membrane-bound PEX5 (green) and PEX14 (red). The brightness of the signal for PEX5, PEX14 and eGFP-SKL appears lower in the case of PEX14(6A). 4×4 µm intakes shown to highlight the details of protein distribution at the peroxisomal membrane. In these intakes brightness and contrast are adjusted to highlight the morphology of the stained proteins. **B)** Images shown in A were used for intensity analysis. The intensity of the signals for the PEX5 and PEX14 staining were measured in peroxisomal regions of interest, indicated by the eGFP-SKL signal. In the scatter plots of intensity of PEX5 (left panel) or PEX14 (middle panel) signals and ratio of intensities of PEX to PEX14 (ratio PEX5/PEX14, right panel), the mean values with standard deviation are indicated for PEX 14 and PEX14(6A) expression as labeled. There is a significant decrease in intensity in case of both proteins for PEX14(6A) compared to a PEX14 expression. The ratio of PEX5 to PEX14 increased in case of the PEX14(6A) compared to the wildtype.

## Discussion

Here, we identified and characterized a novel interaction site of two key components of peroxisomal import, PEX5 and PEX14. The core sequence of the interacting region in PEX14 consists of the hexapeptide IPSWQI located C-terminally to the predicted coiled-coil region, which has the capacity to form PEX14 homo-oligomers ^55^. The PEX14 peptide docks into a cleft formed between the TPR repeats 4 and 5. Notably, this region is distinct from the binding site for the PTS1 peptide of PEX5 cargo proteins but has been suggested to represent a hinge that enables structural rearrangements of the PEX5 TPR supercoil fold upon PTS1 binding ^42^. This important hinge region of the PEX5 TPR domain (residues 470-510), shows distinct conformations in the apo state (more open supercoil of the TPR domain) compared to bound to PTS1 (closed supercoil) ^42^. It is thus tempting to speculate that the binding of the PEX14 peptide in this region may affect the conformation of PEX5 TPR domain and thereby modulate the binding of PTS1. Consistent with this, PEX14 binds with higher affinity to the cargo-free compared to the PTS1-bound PEX5 TPR domain. As seen in structures of PEX5 TPR with PTS1 cargo, *i.e.*, an SKL-peptide ^51,42^ or the SCP2 cargo protein REF, these groove is widened, rationalizing the reduced binding affinity of PEX14 to cargo-loaded PEX5.

For human cells, but also for other organisms, it was generally believed that PEX14/PEX5 interactions are essentially based on the high affinity binding of the conserved N-terminal domain of PEX14 with multiple (di)aromatic peptide repeats (WxxxF-motifs), which are distributed throughout the unstructured N-terminal half of PEX5 ^28, 29, 36, 37, 55^. As an exception, in the yeast *Saccharomyces cerevisiae*, two-hybrid analyses revealed an additional binding PEX14/PEX5 interface that involves the C-terminal 100 amino acids of Pex14 ^27, 56^. Noteworthy, the *S. cerevisiae* Pex14 contains three copies of an IPSWQI-related sequence within its C-terminal Pex5-interacting region. A closer look at the PEX14 sequences from different species revealed that all animal and fungal PEX14 sequences possess a consensus motif with the signature [IL]-P-[SA]-W-[QK]-[ILQNK], either as a single motif or as up to three repeats close to their C-termini (Supplementary Fig. 9). Remarkably, other organisms like plants and Protista lack such motifs in their PEX14 sequences.

We found that disruption of the C-terminal PEX5/PEX14 interface by mutagenesis in human cells causes partial import defects with the most drastic effect on PTS1 proteins. In line with our observations, expression of C-terminally truncated forms of PEX14 lacking the IPSWQI motifs could rescue, at least partially import defects of PEX14 deficient Chinese hamster ovary and *Hansenula polymorpha pex14Δ* cells but resulted in a partial mislocalization of peroxisomal enzymes ^55, 57^. In mammals, a truncated rat PEX14, comprising amino acids 21 to 200, i.e. lacking the IPSWQI sequence, only partially restored the import of PTS1 and PTS2 marker proteins and failed completely to complement the import of catalase that contains a PTS1 with weaker PEX5 affinity ^58^. Instead, the longer PEX14 fragment 21-260 did complement the import import defect for catalase ^55^. In light of our results, this observation might be explained by the fact that the additional fragment from 200-260 contains the novel PEX5 binding site. In Bakeŕs yeast, truncation of the C-terminal region of Pex14, which contains three copies of this motif had a severe defect on the import of both PTS1- and PTS2-import pathways ^27, 56^. However, it is not clear whether the observed strong impact on matrix protein import in *S. cerevisiae* depends on the deletion of the potential Pex5 binding motifs or of other functional elements that reside in the truncated C-terminal region of yeast Pex14, including a Pex7 binding site ^27^.

What could be the function of the C-terminal interface between PEX14 and PEX5 in fungal and human cells? PEX14 has been characterized as a multitasking key player in the processing of the import receptor at the peroxisomal membrane. PEX14 may contribute a primary docking site for cargo-loaded receptors in yeast and humans ^59, 60^, be a constituent of transient translocation pores for PTS1- and PTS2 proteins in yeast ^30, 61^, and contribute to cargo release in human cells ^62^. Expression of PEX5(2A) or PEX14(6A) in PEX5- or PEX14-deficient cell lines at least partially restores peroxisomal protein import arguing against an essential role of the binding interface for the formation of the PTS1-protein import channel. On first sight, the increase of PEX5 at the peroxisomal membrane in comparison to the decreased amount of PEX14(6A) at the membrane and the observation that PEX14 binds with higher affinity to the cargo-free ‘open’ TPR-domain than to the cargo-containing ‘closed’ conformation might suggest that PEX14-binding to the PEX5 TPR domain occurs upon or after cargo-release and argues against a critical role of the binding interface for docking. However, in comparison to wild-type cells, the amount of PEX5 at the membrane is significantly decreased upon expression of PEX14(6A) in PEX14-deficient cell lines, which would support a contribution of the newly identified binding interface in receptor docking or a contribution to pore formation (see below). Recently, a fishing rod-like architecture has been reported for the yeast Pex14/Pex17 complex that suggests a pivotal role of Pex14p in recruitment of Pex5p/cargo complexes ^39^. The rod-like domains position the flexible C-terminal tail of Pex14p and thus the newly identified binding interface far into the cytosol. This could increase the concentration of PEX5 to trigger pore formation as proposed by the transient pore model in yeast ^2^.

Another observation is that the steady state-level of PEX5(2A) is reduced in various PEX5-deficient cell lines, while the amount associated at the peroxisomal membrane is not so much affected and more importantly, the relative amount bound to PEX14 is even increased. This reflects a phenotype that is characteristic for Zellweger patient cell lines defective in receptor recycling. These cell lines show an accumulation of PEX5 at the peroxisomes coupled with a strikingly lower steady-state levels in whole cell lysates ^19^. Accordingly, the reduced efficiency of matrix protein, accumulation of the import receptor at the membrane, and decreased presence in the cytosol can be reconciled with a function of the PEX5/PEX14 interface during or after cargo release from the PTS1 receptor, maybe in receptor recycling. It can be assumed that the accumulation of PEX5 at the peroxisomal membrane either results in ubiquitination of the receptor, as seen for AAA-complex deficient cells ^63^, which might be removed by proteasomal degradation or the whole organelle can be eliminated by ubiquitin-induced pexophagy ^64^ Along this line, it has recently been shown that the loss of PEX13 caused an accumulation of ubiquitinated PEX5 on peroxisomes and an increase in peroxisome-dependent reactive oxygen species that coalesce to induce pexophagy ^58^. It seems likely that both quality control pathways are activated thereby providing rationale for the lower steady-state level of receptor as well as the reduced number of peroxisomes.

The observation that PEX14 binds with higher affinity to the cargo-free ‘open’ TPR-domain than to the cargo-containing ‘closed’ conformation suggests that PEX14-binding to the PEX5 TPR domain occurs upon or after cargo-release. Cargo-release could be triggered by other domains of PEX14, other proteins or environmental changes, like engulfment of the complex by lipids. For instance, binding of the seventh and eighth WxxxF-motifs of human PEX5 by the N-terminal domain of PEX14 was shown to induce PEX5/catalase dissociation ^62^. In yeast cells, Pex8, a peroxin that has not yet been identified in mammalian cells, seems to be involved in cargo dislocation ^65^. Thus, a competitive interaction between the PTS1 sequences of Pex8 and cargo proteins has been considered ^66^. With respect to the newly identified PEX5/PEX14 interface, it seems possible that blocking the hinge region of PEX5 hinders already released PTS1 proteins from re-binding to the TPR-domain. In this model, mutagenesis of the binding sites might result in slower release of cargo and consequently, less-efficient processing of the receptor by the export machinery and finally, removal of the receptor by proteasomal degradation.

Recently, it has been shown that the YG-rich region in PEX13 establishes the formation of a phase-separated condensate that resembles the hydrogel-like meshwork of the nuclear pore ^33, 34^. It has also been shown that PEX5 and PEX14 can simultaneously partition in these condensates together with PEX13, suggesting a mechanism for protein translocation where cargo-loaded PEX5 crosses the membrane through these condensates. While molecular and structural details of these condensates have yet to be characterized, it is well-established that multivalent and weak interactions between components are required to establish such a condensate ^67^. Strikingly, all key peroxins implicated in protein translocation (PEX5, PEX13, PEX14) harbor extended intrinsically disordered regions with peptide motifs that mediate protein-protein interactions with globular domains ^15^. This includes the (di)aromatic peptide motifs in PEX5 that are recognized by the PEX14 NTD and the PEX13 SH3 domain. The newly identified interaction between the PEX14 IPSWQI peptide motif and the PEX5 TPR domain adds another protein-protein interaction to this list. Although this interaction is rather of low affinity, the oligomerization of PEX14 via the coiled-coil domain, the localization and affinity of PEX5, PEX13 and PEX14 at the peroxisomal membrane, and the high local concentration ^34^ in the phase-separated state, argue that these weak interactions are nevertheless highly relevant.

In this respect, the novel PEX14/PEX5 interaction may contribute to all aspects of protein translocation: increasing PEX5 concentration at the membrane, simultaneous partitioning with PEX13 in the condensate and release of PTS1 cargo in the lumen. Future biophysical and structural studies for example, using NMR-methods, should help to unravel the underlying molecular mechanisms. In any case, the recent evidence that phase-separation of key peroxins may rationalize important features of the transient pore model and the role of protein-protein interactions between the components, will help to unravel mechanisms of peroxisomal protein translocation.

## Methods

### Plasmids

Bicistronic expression vectors coding for PEX5L or PEX14, as well as GFP-SKL were described in ^68^. To substitute the PEX14 sequence IPSWQI by six alanine’s, QuikChange (Agilent Technologies, Germany) mutagenesis of the bicistronic plasmid was carried out with a sense (5’ CAGCCCCGAAGGCCGCTGCAGCGGCGGCCCCAGTCAAGTCACCGT 3’) and the corresponding antisense primer. PEX5 Q491 and L494 were replaced by alanine using the primer 5’ CCATTGACCCTGATGTGGCGTGTGGCGCGGGAGTCCTTTTCAACC 3’ and the corresponding antisense primer.

### Protein expression and purification

PEX14 and PEX5 proteins were expressed and purified from a pETM11 vector in *E. coli* BL21(DE3) cells. PEX14 CTRΔcc comprises residues 225 to 377. The full-length and TPR domain of the long isoform (PEX5L) of human PEX5 comprise residues 1 to 639 and 315 to 639, respectively. *E. coli* BL21(DE3) was transformed with the plasmid of interest. An overnight preculture was diluted twenty times in larger volume medium and incubated at 37 °C till OD_600_=0.6-0.8. Expression from plasmids was induced with 200 μM IPTG at 20 °C for 16 hours. Cell pellets were resuspended and sonicated in 50 mM Tris-HCl pH8, 500 mM NaCl, 10 mM imidazole and 1 mM β-mercaptoethanol. Separation of cell debris was achieved by centrifugation at 15,000 g for 20 min. The supernatant was loaded on Ni-NTA column, followed by subsequent washes with buffers 50 mM Tris-HCl pH8, 500 mM NaCl, 10 mM imidazole, 1 mM β-mercaptoethanol, and 50 mM Tris-HCl pH8, 50 mM NaCl, 10 mM imidazole, 1 mM β-mercaptoethanol. Protein of interest was eluted from the column using buffer containing 250 mM imidazole. Subsequently, overnight incubation was performed with TEV protease at room temperature to remove the His_6_-tag and a second nickel column removed the tag and the protease. The isolated protein was then subjected to gel filtration chromatography on S75 column to achieve higher purity. The IPSWQI peptide (PEX14 residues 236-251, preceded by a His_6_-tag and a TEV cleavage site) used for NMR analysis was also produced recombinantly with the same protocol, with only difference using a final Superdex peptide 10/300 column for purification.

Uniformly isotope labeled samples were obtained using M9 medium supplemented with ^13^C-labeled glucose and/or ^15^NH_4_Cl, respectively. For perdeuterated proteins, M9 medium in D_2_O and [U-^2^H]-D-glucose were used for protein expression. For Leu, Val and Ile methyl labeled PEX5 TPR cells were grown in perdeuterated medium containing U-[^2^H]-D-glucose at 37 °C. At OD_600_=0.5, 100 mg of respective precursors were added in the culture. The precursors used were α-ketobutyric acid ^13^C_4_,3,3-d_2_ for (^1^H-^1^δ methyl)-isoleucine and 2-keto 3-methyl ^13^C-butyric 4-^13^C, 3d for (^1^H-^1^δ methyl) -leucine and (^1^H-^1^γ methyl)-valine labeling. Expression and purification was followed as describe above.

The 18-mer IPSWQI peptide (PEX14 residues 234-251, SAPKI<u>PSWQI</u>PVKSPSPS) fused N-terminally with fluorescein isothiocyanate (FITC) and the PTS1 peptide (YQSKL) used for binding experiments were purchased from Peptide Specialty Laboratories GmbH, Heidelberg, Germany. Initially the powder was dissolved in water to achieve 10 mM concentration. Overnight dialysis against water followed to remove any residual trisfluoroacetic acid (TFA).

### NMR spectroscopy

All NMR experiments were performed in 20 mM phosphate buffer, pH 6.5 and 50 mM NaCl. Backbone chemical shift assignments for PEX14 CTRΔcc (residues 225 to 377) and PEX5 TPR protein (residues 315 to 639) was achieved with standard triple resonance experiments HNCA, HNCACB, HNCO, HNCACO, CBCA(CO)NH ^69^. For the 35 kDa PEX5 TPR domain, TROSY versions of the experiments were used; H(CC)(CO)NH and (H)CC(CO)NH experiments were recorded in addition. NMR spectra were recorded on a Bruker AvanceIII 900 MHz spectrometer equipped with a cryogenic probe. Assignments of Ile, Leu and Val groups for the PEX5 TPR domain were obtained from ^13^C-and ^15^N-edited simultaneous NOESY-HSQC spectra in combination with ^1^H-^13^C HMQC, ^1^H-^15^N TROSY and HCC-FLOPSY ^70^ with a NOESY mixing time of 100 ms, and a FLOPSY mixing time of 22 ms. ^1^H chemical shift assignments of the unlabeled peptides were obtained using ^1^H,^1^H TOCSY, ^1^H,^1^H NOESY, and natural abundance ^1^H-^13^C HSQC 2D spectra recorded on a Bruker Avance III 500 MHz spectrometer equipped with a cryogenic probe. Sample concentrations were 0.8-1 mM. NMRpipe ^71^ was used for processing of all spectra, data analysis and assignments were performed using Sparky ^72^. Backbone assignments were assisted by MARS ^73^.

NMR titrations experiments were performed in NMR buffer with 150 µM ^15^N-^13^C-and ILV-labeled PEX5 TPR domain and with increasing amount of 150, 400, 900 and 1800 µM of unlabeled PEX14 IPSQWI peptide.

Intermolecular NOEs between PEX5 TPR and the unlabeled PEX14 IPSWQI peptide were detected from a ω_1_-filtered/ω_3_-edited 3D NOESY-HSQC experiment with 110 ms mixing time. The spectra were recorded on a Bruker Avance 900 MHz spectrometer equipped with a TXl cryoprobe head. Sample consisted of 400 μM PEX5 TPR and 4 mM unlabeled PEX14 IPSWQI peptide. Chemical shifts and NOEs of the bound conformation of the PEX14 IPSWQI peptide were obtained from 2D transferred ^1^H,^1^H NOESY ^74^ spectra with 250 ms mixing time ^75^. The experiment was recorded on Bruker Avance III 500 MHz spectrometer on sample containing 50 μM unlabeled PEX5 TPR and 2 mM unlabeled IPSWQI peptide.

### Structure calculations

To determine a structural model of the PEX14/PEX5 TPR complex we first calculated the PEX5-bound conformation of a PEX14 peptide for residues 237-245 (KI<u>PSWQI</u>PV) with CYANA 3.30 based on distance-restraints derived from transferred NOEs (Supplementary Table 1, for structural statistics). The structure of the human PEX5 TPR domain (2C0M^42^) and the bound conformation of the PEX14 peptide were docked using 10 unambiguous distance restraints derived from intermolecular NOEs (Supplementary Table 2) and 13 ambiguous interaction restraints as active residues derived from CSPs. PEX5 TPR residues with CSP > 0.05 ppm (Lys472, Glu473, Leu476, Val479, Glu480, Asp482, Thr483, Ser485, Ile486, Lys506, Asp509, Ala513, Ser516,) and the K237, I238, P239, S240, W241, Q242, I243 in the KIPSWQIPV peptide. Residues within 6.5 Å distance to active residues were treated as passive residues. For the docking calculation performed at the HADDOCK2.4 web server^47, 48^, 2000, 400 and 200 structures were calculated, in iterations 0, 1 and for final water refinement, respectively. The minimum cluster size was set to 4. Structural statistics of the final selected ensemble is shown in Supplementary Table 3,4. Using a 5 Å root-mean-square-deviation (RMSD) clustering criterion, one structural cluster was identified for a total of 199 (out of 200 calculated) complex structures. The Best 10 structures were selected on HADDOCK score (Supplementary Table 3). The structures were analyzed using HADDOCK and CAPRI and self-developed analysis scripts ^47–49^.

### MD simulation

To avoid simulation artefacts due to missing residues, structures were rebuilt using Modeller^76^.Then 600 ns MD simulations were calculated using the GROMACS environment ^50, 77, 78^. During the simulations, the temperature was controlled after the equilibration steps to be at 300 K and the density close to that of liquid water (1037 kg/m^3^). To process the simulation data, we applied the gCorrelation method of Lange and Grubmueller ^79^, which is an extension of the GROMACS environment. The gCorrelation uses an information theory background to capture the mutual information between residues, measured from MD simulations. This is an effective measure of the correlation of the motions of residues in a biological system observed during a simulation, and may be considered a very compact, yet highly informative, summary of the simulation. The gCorrelation was calculated for the C-alpha, side chains, and main chains. Here we only show and discuss the results for the former, since it provides a more compact description of the data than the main-chain analysis. We noticed however that C-alpha correlations were missing some correlations, so these were extended with an additional side chain analysis.

### Fluorescent polarization assays

For the determination of each binding curve a n-point titration was performed in the NMR buffer with constant concentration of 20 nM FITC-labeled SAPKI<u>PSWQI</u>PVKSPSPS peptide and increasing concentrations of receptor or premixed receptor:PTS1 peptide complex in 1:1 ratio. Reaction mixtures was then transferred into 96-well Optiplate and measured as triplicates in an EnVision plate reader.

### Peptide substitution analysis

Immobolized peptides were synthesized by the Fmoc method of solid phase peptide chemistry on a Milligen/Biosearch 9050 Pep-synthesizer. Cleavage of protecting groups, as well as removal from the solid phase was accomplished by treating the peptide with 90 % trifluoroacetic acid, 2.5 % phenol, 2.5 % 1,2-ehtanedithiol, 1 % triisopropylsilane (all from Sigma) and 5 % water ^80^. A combinatorial library of the human PEX14 (234-248) substitution variants was generated by the Sports technique according to the manufacturers protocol *(ABIMED, Auto-Spot-Robot ASP 222)* as described previously ^80^. The library was incubated with 200 nM purified His_6_-tagged PEX5L. Bound PEX5L was detected immunologically using monoclonal Anti-His Antibodies (Qiagen).

### Structured Illumination Microscopy (SIM) and Analysis

The PEX5-deficient fibroblast cell line ΔPEX5T was described in ^81^, HEK-293 cell lines and human fibroblast cells were cultured as described previously ^42^. Cells were maintained in DMEM (Sigma) supplied with 4500 mg glucose/L, 110 mg sodium pyruvate/L supplemented with 10% fetal bovine serum, 2 mM glutamine and 1 % penicillin-streptomycin in 8.5 % CO_2_ at 37 °C.

For the complementation analysis, bicistronic vectors coding for a PEX5 or PEX14 variant, as well as eGFP-SKL were transfected with Nucleofector (fibroblasts, Lonza) or XtremeGene (HEK cells, Roche). Cells were fixed 48 h after transfection for 20 min with 3 % Formaldehyde/DBPS and permeabilized with 1 % Triton X-100/DPBS for 5 min at room temperature. For blocking, the cells were incubated with 2 % BSA, 5 % FCS (Sigma) in DPBS for 1 h. For immunostaining of the peroxisomal membrane marker PMP70, a polyclonal PMP70-antiserum (PA1-650, Thermo Fischer, 1:500) and a conjugated secondary antibody Alexafluor 594 (Thermo Scientific) was used. Cells were mounted in Mowiol with DAPI (Sigma).

Fluorescence microscopy was performed using a Zeiss ELYRA PS.1 equipped with a 63x oil immersion objective, which allowed generation of high-resolution images using the Structure Illumination Microscopy (SIM) technology. SIM images were processed using the ZEN2.1 software (Carl Zeiss, Jena). Three to five biological replicates were used for quantitative analysis 48 hours after transfection. Based on the appearance of the eGFP-SKL fluorescence pattern, about 100 cells per replicate were visualized and categorized in complemented or import defected cells. Whereas a complementation was indicated by punctuated staining, import defective cells show a cytosolic GFP-staining.

### STED Microscopy Image Acquisition and Analysis

HEK KO PEX14 deletion cells were maintained in a DMEM culture medium with 4500 mg glucose/liter and 110 mg of sodium pyruvate/liter supplemented with 10% fetal calf serum, 2 mM glutamine, and 1% penicillin-streptomycin. The cells were cultured at 37 °C in 8.5 % CO_2_. To make sure the HEK-293 KO PEX14 cells were completely complemented the cells were transfected 5 days before imagining. Therefore, the cells were transferred to number 1.5 cover slides four days after transfection and one day before fixation for the immunolabeling. The cells were transfected with dual pIRES2 expression plasmids containing the eGFP-SKL marker protein and PEX14 or the modified PEX14(6A). Transfection was carried out using Lipofectamine 2000 transfection reagent (Invitrogen). The immunolabeling was performed as described before ^54^.

STimulated Emission Depletion (STED) microscopy image acquisition was performed using frame-mode on the Abberior system, as described elsewhere ^54, 82^. Due to the frame-based acquisition, some drift was also likely present and so this effect was measured through alignment of the confocal acquired channels which accompany each STED microscopy image acquisition. Once measured, the appropriate correction was applied to both the confocal and STED microscopy images. Next, each peroxisome was detected through using a maxima finding algorithm applied to the confocal channel (eGFP-SKL) acquired first. At each detection point a circular region was analyzed of diameter 21 pixels. In this region the intensity of each pixel was measured and the mean intensity calculated.

For each peroxisome region the intensity in the first channel image and the two STED microscopy images were measured. Additionally, the ratio between the measured mean intensities was calculated for each region to provide a normalised ratio for each measurement.

We plotted the data from three independent experiments with 10 STED microscopy images each. The samples for each condition were processed and imaged in parallel on the same day to ensure constant conditions of labelling and STED microscopy image acquisition.

## Competing interests

The authors declare that they have no competing interests.

## Accession codes

Structural coordinates and restraints for the calculation of the PEX14 peptide/PEX5 TPR complex are deposited at the PDB and BMRB with the accession codes 8Q4C and 34842. NMR chemical shifts for the different PEX5 and PEX14 constructs are deposited at BMRB with the accession codes 52048, 52054 and 52055.

## Author contributions

L.E. performed protein expression, biophysical experiments, NMR and structural analysis. D.G., J.K. and S.G. contributed to protein expression and NMR analysis. K.R. executed the creation and screening of the CRISPR/Cas9 deletion cell lines, STED high resolution imaging and data analysis. J.S. performed SIM microscopy imaging and the analysis of import efficiency. D.W. wrote the algorithm for the intensity analysis of the STED images. P.H. designed the strategy for creation of the CRISPR/Cas 9 deletion cell lines. V.B. and K.F.W. provided SIM instrumentation and help with data analysis. M.J. provided peptide blots. F. M. performed and analyzed MD simulations. M.S., W.S., R.E. and C.E. conceived the project; L.E, W.S., R.E. and M.S. wrote the manuscript. All authors discussed the project and approved the manuscript.

## Acknowledgements

We thank Sam Asami and Gerd Gemmecker, BNMRZ, for help with NMR experiments, and Jacob Anglister, Weizmann, Israel, for discussions. We thank the Wolfson Imaging Centre Oxford for microscope facility support with data acquisition and analysis, as well as the Microverse Imaging Center (Aurélie Jost; Patrick Then) for providing microscope facility support.

## Funding

This work was supported by grants of the Deutsche Forschungsgemeinschaft (DFG): FOR1905 (project number 219314758) to C.E. (EG325/1-2), M.S. (Sa823/7, Sa823/11), and R.E. (Er178/6, Er178/7). C.E. acknowledges further funding by the DFG (Germanýs Excellence Strategy – EXC 2051 – Project-ID 390713860; project number 316213987 – SFB 1278; INST 275_405_1; INST 1757/25-1 FUGG funding), by the State of Thuringia (TMWWDG, project ThIMEDOP), by the MRC (Grant No. MC_UU_12010/unit programs G0902418 and MC_UU_12025), by the Wellcome Trust (Grant No. 104924/14/Z/14 and Strategic Award 091911 (Micron)), MRC/BBSRC/EPSRC (Grant No. MR/K01577X/1), by the Wolfson Foundation (for initial funding of the Wolfson Imaging Centre Oxford), and integration into the Leibniz Center for Photonics in Infection Research (LPI, initiated by Leibniz-IPHT, Leibniz-HKI, UKJ and FSU Jena and part of the BMBF national roadmap for research infrastructures). K.F.W is supported by DFG grants WI 2111/6; WI 2111/8) and EXC2033 - project number 390677874 – RESOLV. SIM microscopy was funded by the DFG and the State Government of North Rhine-Westphalia (INST 213/840-1 FUGG).

## Notes

### Competing Interest Statement

The authors have declared no competing interest.

