## Supporting Information for "A novel PEX14/PEX5 interface links peroxisomal protein import and receptor recycling"

##### Contents

***Supplementary Tables***


| | | |
|---|---|---|
| Suppl. Table 1 | Structural statistics for the PEX5-bound PEX14 IPSWQI peptide conformation | 2 |
| Suppl. Table 2 | Intermolecular NOEs PEX14 IPSWQI / PEX5 TPR | 3 |
| Suppl. Table 3 | Structural statistics for the 10 best structures from semi-rigid body docking of the PEX14 IPSWQI/PEX5 TPR complex | 4 |
| Suppl. Table 4 | HADDOCK score and energies for the PEX14 IPSWQI /PEX5 TPR complex | 5 |


***Supplementary Figures***


| | | |
|---|---|---|
| Suppl. Figure 1 | NMR spectra of PEX14 CTR∆cc free and bound to full length PEX5 | 6 |
| Suppl. Figure 2 | Conformation of the PEX5-bound PEX14 IPSWQI peptide | 7 |
| Suppl. Figure 3 | Intermolecular NOEs PEX14 IPSWQI /PEX5 TPR | 8 |
| Suppl. Figure 4 | Structural analysis of the 10 best HADDOCK structures with details on the PEX5 TPR - PEX14 KIPSWQIPV interface | 9 |
| Suppl. Figure 5 | Analysis of MD trajectories from the complex PEX5 TPR with PEX14 IPSWQI peptide | 10 |
| Suppl. Figure 6 | Mutational analysis of the interaction of an PEX14 IPSWQI-containing peptide with PEX5 | 11 |
| Suppl. Figure 7 | Functional complementation of PEX5-deficient fibroblasts | 12 |
| Suppl. Figure 8 | PEX5L$^{mut}$ shows decreased stability | 13 |
| Suppl. Figure 9 | The PEX5-binding motif of PEX14 is conserved in animals and fungi | 14 |

#### Supplementary Table 1

**Structural statistics the PEX14 IPSWQI peptide bound to PEX5**

| | |
|---|---|
| **NMR restraints** | |
| Distance restraints | |
| Total # NOEs | 40 |
| Inter-residue | |
| Sequential (\|*i* – *j*\| = 1) | 35 |
| Medium-range (\|*i* – *j*\| < 4) | 5 |
| Long-range (\|*i* – *j*\| > 5) | 0 |
| **Restraint statistics** | |
| Violations (mean ± s.d.) | |
| Distance restraints (Å) | 0.003 ± 0.01 |
| Max. distance restraint violation (Å) | 0.32 |
| **Strutural quality** | |
| Deviations from idealized geometry | |
| Bond lengths (Å) | 0.001 ± 0.00 |
| Bond angles (º) | 0.2 ± 0.07 |
| **Ramachandran plot (%)** | |
| Residues in most favored regions | 49 |
| Residues in additionally allowed regions | 51 |
| Residues in generously allowed regions | 0 |
| Residues in disallowed regions | 0 |
| **Precision,** coordinate r.m.s. deviation (Å)* | |
| Backbone | 0.67 ± 0.27 |
| Heavy atoms | 1.17 ± 0.34 |

*Pairwise coordinate r.m.s. deviation was calculated for 20 refined structures for the IPSWQI core motif, PEX14 residues 238-243.

#### Supplementary Table 2

##### Intermolecular NOEs PEX14 IPSWQIP / PEX5 TPR

| PEX5 TPR | PEX14 KIPSWQIV peptide |
|---|---|
| Leu476 HD## | K237HB# / I238HB / P239 HG# / Q242 HB# / I243HB / P244 HB# |
| Leu476 HD## | W241 HE3 / HD1 / HZ3 / HH2 / HZ2 |
| Val479 HG1# | I238 HB / P239 HG# / Q242 HB#/ Q242 HG# / I243 HB / P244 HB# |
| Val479 HG1# | I238 HD1# / I243 HD1# |
| Val479 HG1# | I238 HG# / I243 HG# |
| Ile486 HD1# | I238 HG# / I243 HG# |
| Ile486 HD1# | I238 HG# / I243HG# / V245 HG# |
| Ile486 HD1# | I238 HA / P244 HA |
| Ile486 HD1# | I238 HB / Q242 HB# / I243HB / P244 HB# |
| Ile486 HD1# | P239 HG# / Q242 HG# |

#### Supplementary Table 3

**Structural statistics for the 10 best structures from semi-rigid body docking of the PEX14 IPSWQI/PEX5 TPR complex**

| | |
|---|---|
| **NMR restraints** | |
| Distance restraints | |
| # intermolecular NOEs | 10 |
| # CSP-derived distance restraints | 13 (PEX5) + 7 (PEX14) |
| **Structural statistics** | |
| Violations (mean ± s.d.) | |
| Distance restraints (Å) | 0.19 ± 0.74 |
| Max. distance restraint violation (Å) | 0.26 |
| Deviations from idealized geometry | |
| Bond lengths (Å) | 0.003 ± 0.000 |
| Bond angles (°) | 0.400 ± 0.005 |
| **Ramachandran plot statistics (%)** | |
| Residues in most favored regions | 94.1% |
| Residues in additionally allowed regions | 3.3 % |
| Residues in generously allowed regions | 0 % |
| Residues in disallowed regions | 2.5 % |

#### Supplementary Table 4

**HADDOCK score and energies for the PEX14 IPSWQI /PEX5 TPR complex.**

| HADDOCK statistics | | |
|---|---|---|
| **Parameter** | **Best 10** | **Best** |
| HADDOCK score | -73.6 ± 2.9 | -79.6 |
| RMSD[a] (Å) | 0.7 ± 0.1 | 0 |
| i-RMSD[b] (Å) | 0.47 ± 0.22 | 0 |
| l-RMSD[c] (Å) | 2.1 ± 1.4 | 0 |
| E-vdW[d] (kcal/mol) | -44.9 ± 4 | -45.7 |
| E-elec[e] (kcal/mol) | -164 ± 23.1 | -165.8 |
| E-vio[f] (kcal/mol) | 33.9 ± 19.0 | 11.9 |
| E-desolv[g] (kcal/mol) | 0.8 ± 1.9 | -1.9 |
| BSA[h] (Å) | 1148.9 ± 37.9 | 1146 |
| Fnat[i] | 0.81 ± 0.11 | 1 |

[a] root mean square deviation (of pairwise alignment to the best structure)
[b] interface root mean square deviation
[c] ligand root mean square deviation
[d] van der Waals contribution to intermolecular energies
[e] electrostatic contribution to intermolecular energies
[f] restraint violation energy
[g] desolvation energy
[h] buried surface area
[i] fraction of native contacts

### Supplementary Figures

#### Supplementary Figure 1

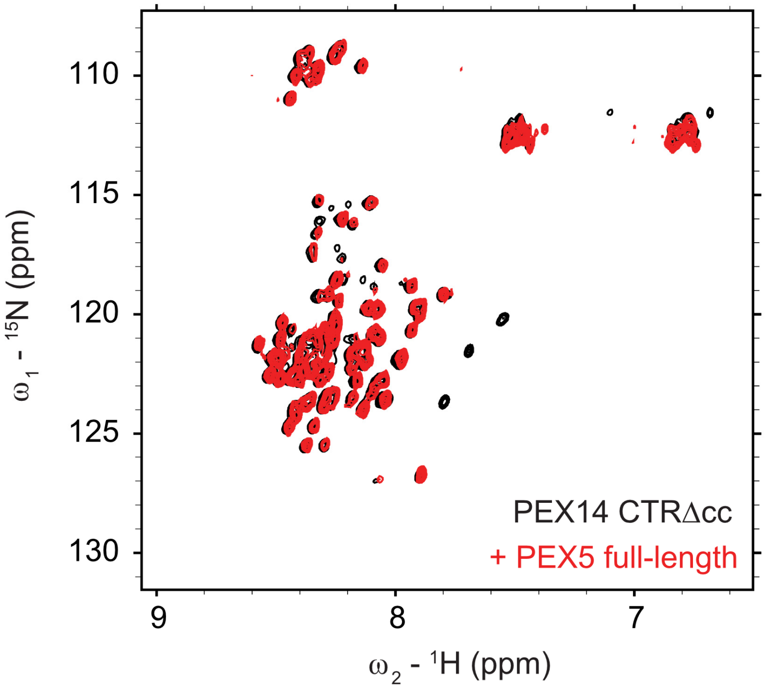


**Supplementary Figure 1: NMR spectra of PEX14 CTRΔcc free and bound to full length PEX5.** $^{1}$H,$^{15}$N HSQC spectra of 100 μM PEX14 CTRΔcc free (black) and in presence of an equimolar amount of full length PEX5 (red).

#### Supplementary Figure 2

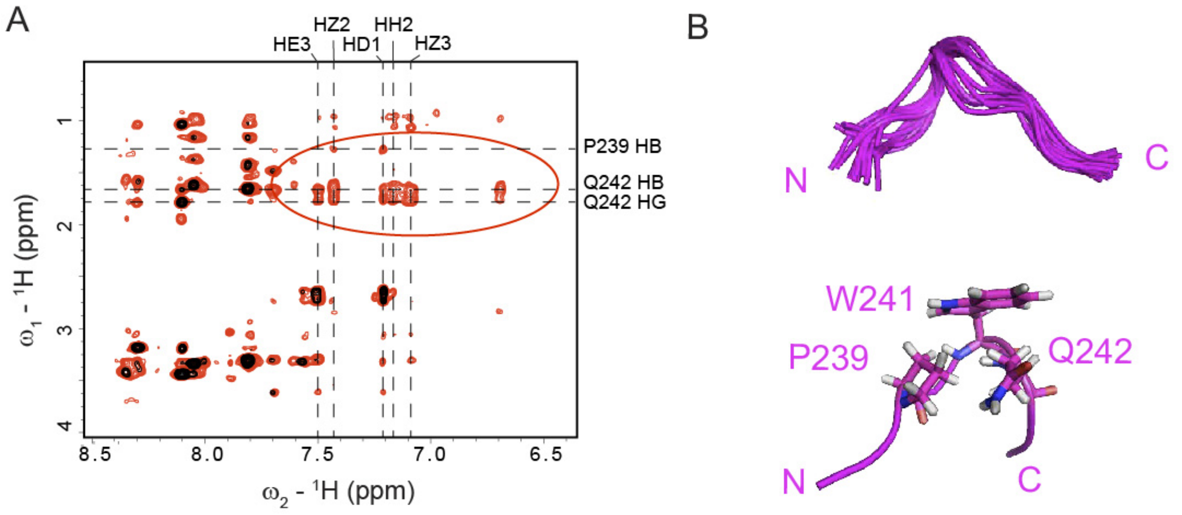


**Supplementary Figure 2: Conformation of the PEX5-bound PEX14 IPSWQI peptide**
**A)** Transferred NOE experiment showing the amide – aliphatic region. The black spectrum was obtained from a 1 mM solution of the unlabeled PEX14 IPSWQI peptide; while in red 50 μM of unlabeled PEX5 TPR is added. The intense additional peaks highlighted in circle correspond to transfer NOEs from the bound conformation of the peptide to the free. Vertical dashed lines report the resonances of the aromatic protons of the tryptophan (W241), while horizontal lines represent proline (P239) and glutamine (Q242) protons, which show intramolecular NOEs towards the tryptophan W241. **B)** NOE-based structural ensemble of the central region of the peptide. The 10 lowest energy structures are shown (top), important side chains are shown as sticks for the lowest energy structure (bottom).

#### Supplementary Figure 3

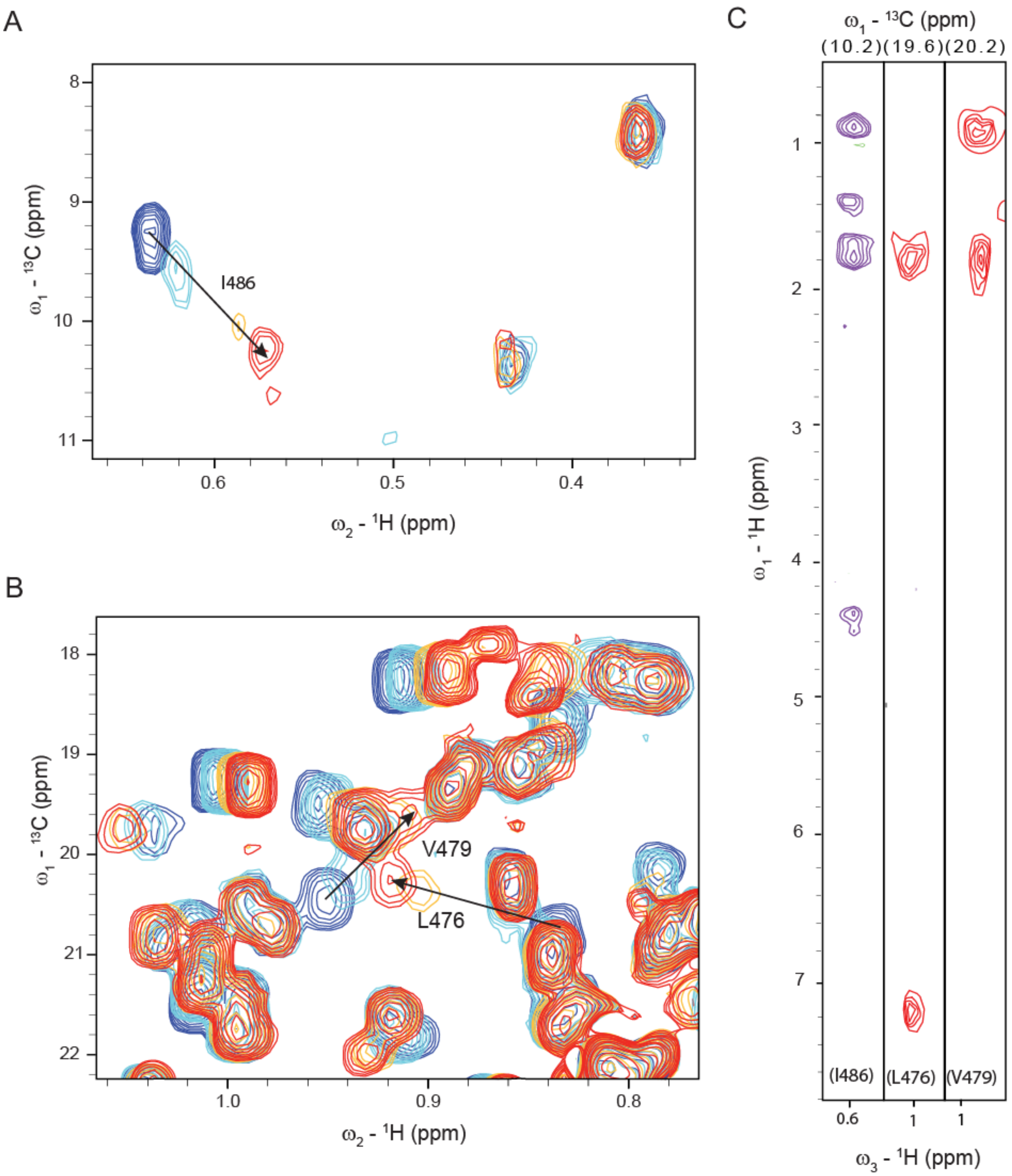


**Supplementary Figure 3. Intermolecular NOEs between PEX5 TPR and the PEX14 IPSWQI peptide. A)** $^{1}H,^{13}C$ HMQC spectra overlay of PEX5 isoleucine δ1 methyls at 100 µM protein concentration (dark blue) with 100 µM (light blue), 300 µM (orange) and 1000 µM (red) unlabeled IPSWQI peptide. Isoleucine I486 δ1-methyl showing the largest chemical shift perturbation and intermolecular NOEs is displayed. **B)** Same titration as in A zoomed in the Leu γ1- and Ile δ1- methyls region. Methyls exhibiting NOEs with IPSWQI peptide are marked. **C)** $\omega_1$-filtered/$\omega_3$-edited NOESY strips displaying intermolecular NOEs from Pex5 I486 δ1-methyl (purple), L476 γ1-methyl and V479 γ1-methyls (red) to the IPSWQI peptide.

**Supplementary Figure 4**

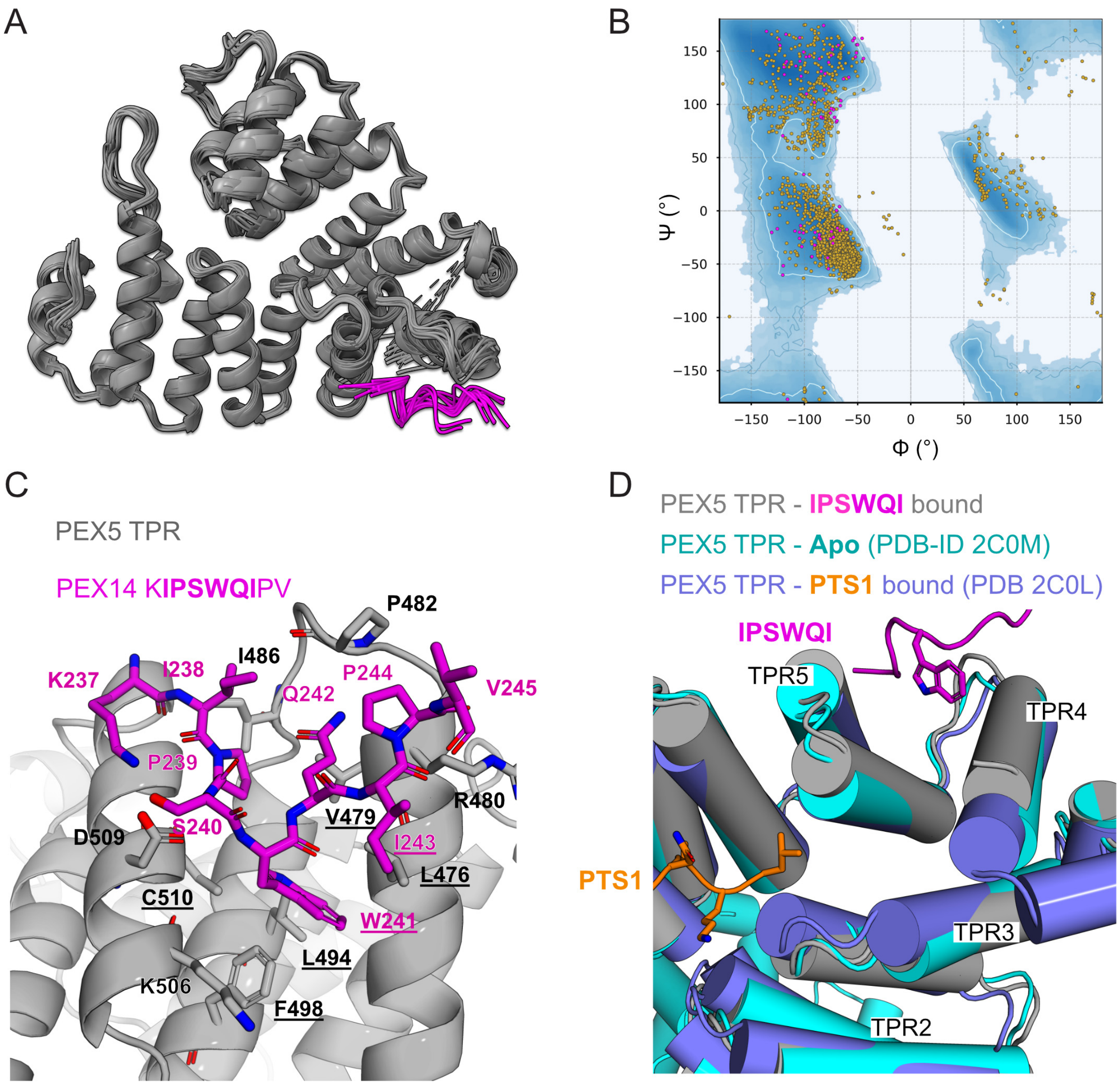


**Supplementary Figure 4**: **Structural analysis of the 10 best HADDOCK structures with details on the PEX5 TPR - PEX14 KIPSWQIPV interface. A)** Overlay of the 10 best HADDOCK structures. The PEX14 IPSWQI peptide of the best structure is shown in magenta while others are shown in light pink. **B)** Ramachandran pot of the best 10 complex structures. Backbone angles from PEX5 TPR are shown in orange and from PEX14 are shown in magenta. **C)** Binding of the PEX14 peptide is mainly mediated by hydrophobic contacts, which is centered by the Tryptophane W241. The hydrophobic pocket on PEX5 is formed L476, V479, P482, I486, L494, F498, C510 and the aliphatic sidechains of R480 and K506, accommodating the hydrophobic residues of the PEX14 KIPSWQIPV peptide. **D)** Overlay of the PEX5 TPR apo, the PTS1 bound, and the WQI bound form shows a structural rearrangement of the TPR segments upon PTS1 or WQI binding in opposite directions.

#### Supplementary Figure 5

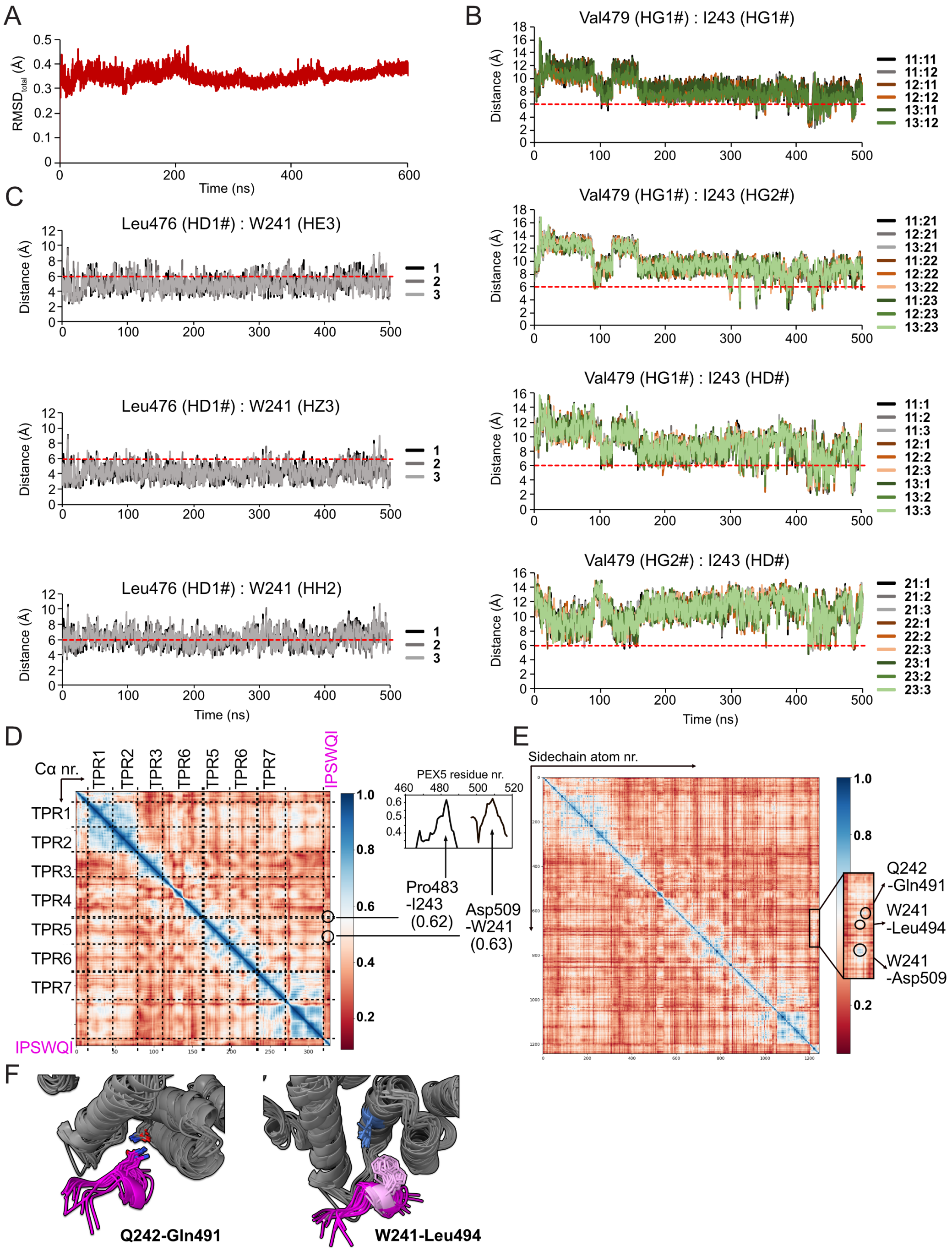


**Supplementary Figure 5. Analysis of MD trajectories from the complex PEX5 TPR with PEX14 IPSWQI peptide. A)** The RMSD was calculated on the whole protein complex (excluding protons). The stable RMSDs of each frame with respect to the initial structure indicates a stable complex over the full simulation time. **B)** Over the simulation time of 500 ns traced distances between selected Val479 HG## and I243HG## as well as I243HD# show distances below the threshold for observable NOE signals of 6 Å after 300-400 ns simulation time. **C)** Traced distanced between Leu476 HD1# and W241 He3, Hz3 and Hh2 protons are always in a good observable distance of 2 to 6 Å. **D)** Ca atom gCorrelation matrix (over the full simulation time) of the complex shows correlation of Pro483 with I243 and Asp509 with W241. **E)** Sidechain gCorrelation matix of the PEX5 TPR – PEX14 IPSWQI complex reveals correlations between Gln242 and Gln491, Trp242 and Leu494, as well as Trp241 and Asp509. Those interactions are visualized in panel **F.**

**Supplementary Figure 6**

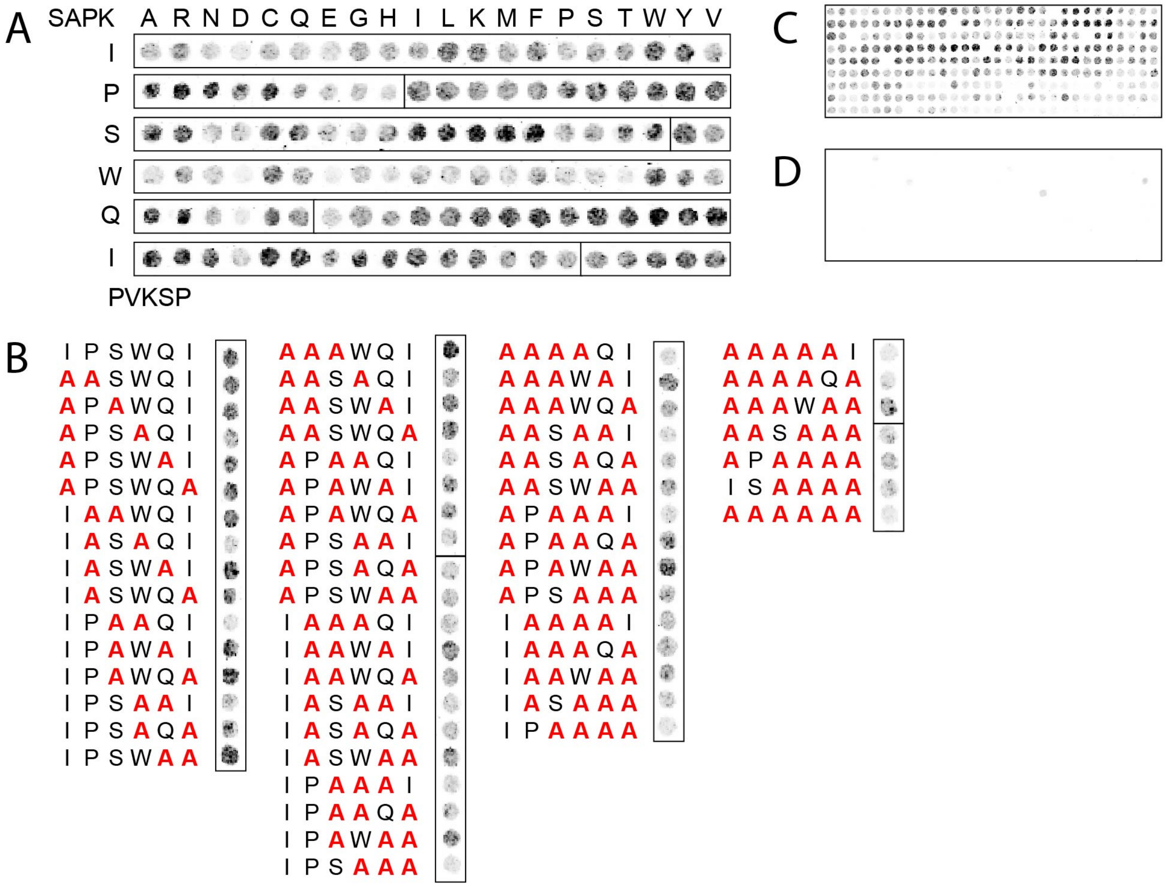

**Supplementary Figure 6**: **Mutational analysis of the interaction of an PEX14 IPSWQI-containing peptide with PEX5.** Peptides comprising systematic variations of the IPSWQI motif in PEX14 (residues 234-248; SAPKIPSWQIPVKSP) were synthesized on cellulose membranes and incubated with purified $His_6$-PEX5L. Bound PEX5 was visualized by immunodetection with polyclonal PEX5-antibodies. Spots with reduced intensities represent peptides with reduced binding affinities for PEX5L. **A)** Substitution analyses were performed with 15-mer peptides, where each amino acid of the PEX5-binding IPSWQI sequence was replaced with any of the 20 amino acids or **B)** two, three, four or five amino acids of the core sequence were changed to alanines. To demonstrate the specificity of labelling, the cellulose membrane with all immobilized peptides was incubated **C)** with and **D)** without PEX5 followed by immunodetection.

#### Supplementary Figure 7

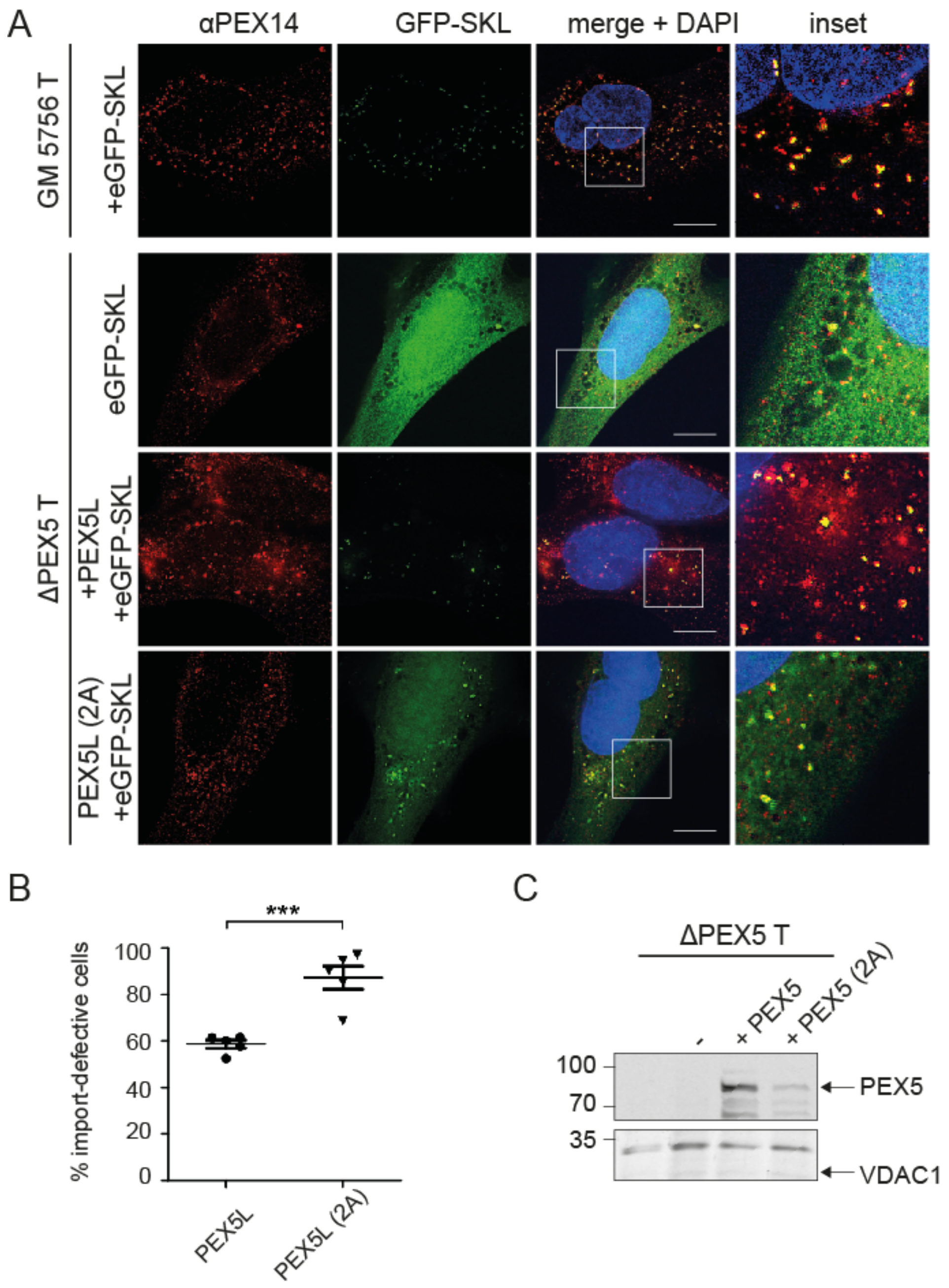


**Supplementary Figure 7: Functional complementation of PEX5-deficient fibroblasts** using wildtype PEX5 and a PEX5 Q491A L494A mutant. Both constructs were expressed in PEX5-deficient fibroblasts (PEX5-001T) from bicistronic expression vectors coding for the full-length PEX5 as indicated and a eGFP-SKL fusion protein as marker for PTS1 matrix protein import. **A)** Peroxisomes were visualized with anti-PEX14 antibodies (red channel) and images were obtained via SR-SIM. Scale bars: 10 µm. **B)** Quantification of the restoration of peroxisomal PTS1-import. For each experiment, 100 cells were categorized by the localization of the PTS1-import marker eGFP-SKL. Cells with a diffuse cytosolic staining of eGFP-SKL show at least a partial import defect. Values were obtained from five independent experiments. Error bars are shown as mean with SD. For statistical analysis, values were tested for Gaussian distribution using Kolmogorov-Smirnov-test and significance was determined by one-way analysis of variance Anova (*p<0,05; **p<0,01; ***p<0,001). **C)** The expression level and stability of the PEX5 variants was checked by immunoblot analysis using specific polyclonal PEX5 antibodies. An empty vector control (pIRES2) shows the specificity of the antibody.

#### Supplementary Figure 8

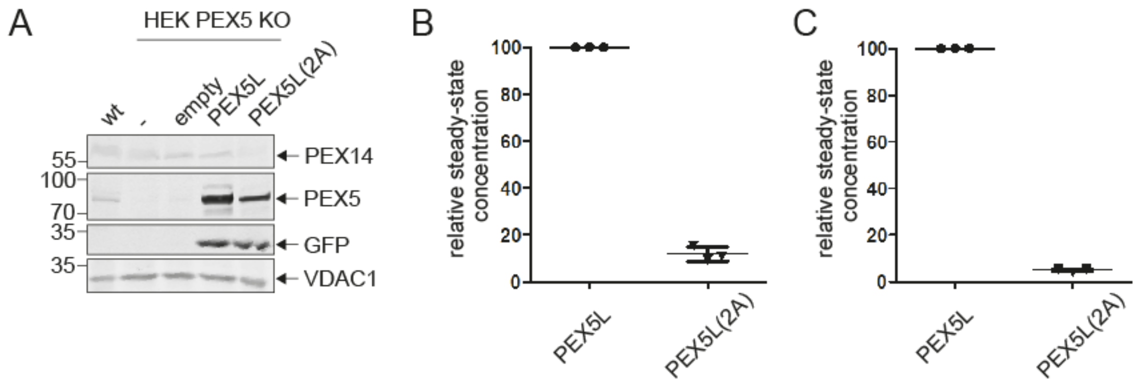


**Supplementary Figure 8: PEX5L(2A) shows decreased stability. A)** Whole cell lysates were prepared from HEK-293 (wt), HEK-293 PEX5 Knockout (HEK PEX5 KO) cells, and PEX5 KO cells, transfected with an empty bicistronic control vector encoding eGFP-SKL alone (empty) or, in addition one of the indicated PEX5 variants wild-type PEX5L and mutant PEX5L(2A). Equal amounts were analyzed by immunoblotting of PEX14, PEX5, GFP and VDAC1 as a loading control. **B,C)** Relative PEX5L and PEX5L$^{mut}$ steady-state concentrations were obtained from the densitometric analysis of signal intensities in cell lysates from HEK PEX5 KO cells **(B)** and PEX5-deficient fibroblasts derived from Zellweger patients **(C).** The signal intensity of VDAC1 immunostaining was used as an internal standard.

#### Supplementary Figure 9

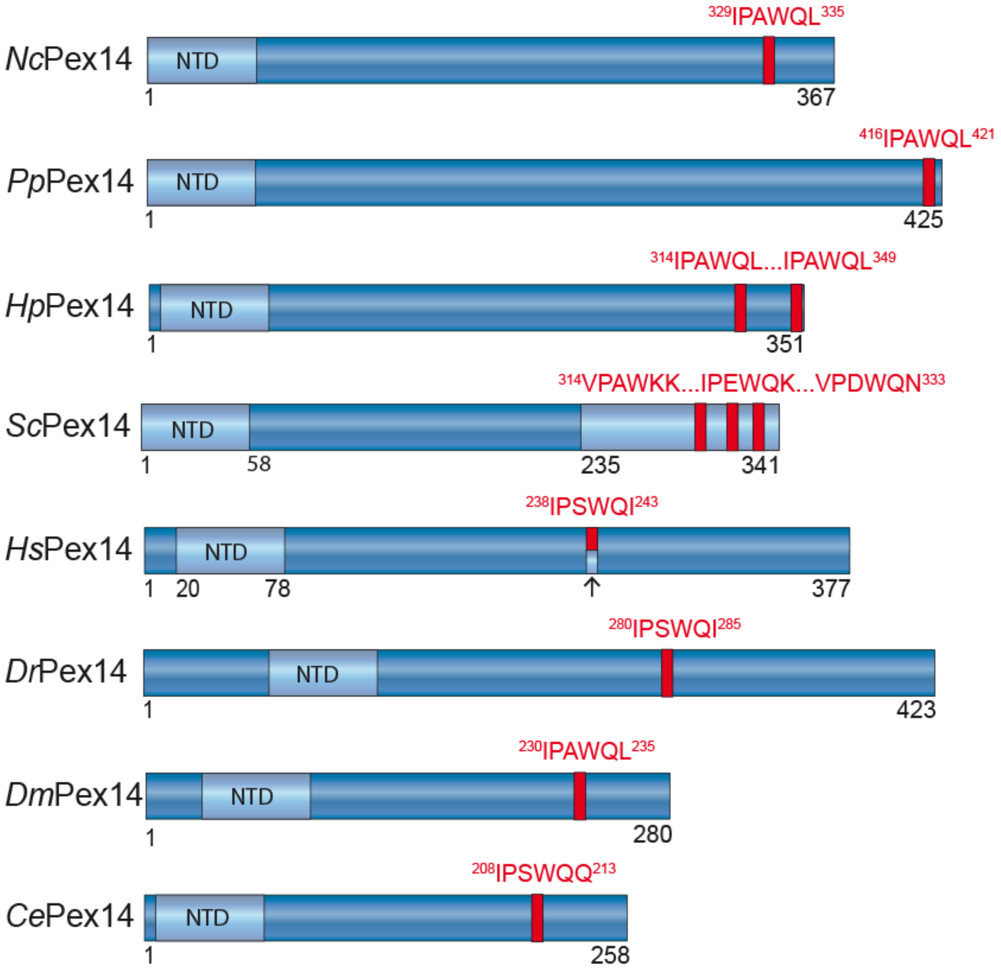


**Supplementary Figure 9: The PEX5-binding motif of PEX14 is conserved in animals and fungi.** Representative PEX14 sequences are indicated as blue tubes. Numbers indicate position of amino acids at the boundaries of relevant structural and functional regions. The [I,V]-P-[X]-W-[Q,K]-[L,I, Q, K, N] sequences (indicated as red bars and sequences above) are located in one to three copies between coiled-coil regions and extreme C-termini of the indicated PEX14 sequences. Experimentally identified PEX5-interacting regions within ScPEX14 and HsPEX14 are shown in light blue. Abbreviations used for organisms are *Nc, Neurospora crassa; Pp, Pichia pastoris; Hp, Hansenula polymorpha; Sc, Saccharomyces cerevisiae; Hs, Homo sapiens; Dr, Danio regio; Ce, Caenorhabditis elegans; Dm, Drosophila melanogaster.*
